# Identifying the drivers of computationally detected correlated evolution among sites under antibiotic selection

**DOI:** 10.1101/474536

**Authors:** Jonathan Dench, Aaron Hinz, Stéphane Aris-Brosou, Rees Kassen

**Affiliations:** Department of Biology, University of Ottawa, Ottawa, ON K1N 6N5, Canada; Department of Mathematics and Statistics, University of Ottawa, Ottawa, ON K1N 6N5, Canada

**Keywords:** Correlated evolution, epistasis, antibiotic resistance, *Pseudomonas aeruginosa*, computational methods, experimental validation

## Abstract

The ultimate causes of correlated evolution among sites in a genome remain difficult to tease apart. To address this problem directly, we performed a high-throughput search for correlated evolution among sites associated with resistance to a fluoroquinolone antibiotic using whole genome data from clinical strains of *Pseudomonas aeruginosa*, before validating our computational predictions experimentally. We show that for at least two sites, this correlation is underlain by epistasis. Our analysis also revealed eight additional pairs of synonymous substitutions displaying correlated evolution underlain by physical linkage, rather than selection associated with antibiotic resistance. Our results provide direct evidence that both epistasis and physical linkage among sites can drive the correlated evolution identified by high throughput computational tools. In other words, the observation of correlated evolution is not by itself sufficient evidence to guarantee that the sites in question are epistatic; such a claim requires additional evidence, ideally coming from direct estimates of epistasis, based on experimental evidence.

## 1 INTRODUCTION

Adaptive evolution often involves changes to multiple characters (or traits) in concert, a process called correlated evolution. Such coordinated changes arise either because of physical linkage in the genome, or from strong selection that generates an association between distinct genomic sites. The analysis of correlated evolution among traits has a long history in quantitative genetics (Falconer, 1960; Cheverud, 1984; Lynch and Walsh, 1998), molecular biology (Goh *et al.*, 2000; Callahan *et al.*, 2011), life history evolution (Baer and Lynch, 2003; Sexton *et al.*, 2009; Kelley *et al.*, 2013), and comparative biology (Pagel, 1994; Shimizu *et al.*, 2014). A comparable effort at the genomic level remains a daunting task because the number of potentially interacting sites (*i.e.* nucleotides) can be overwhelmingly large, making it difficult to distinguish genuine instances of correlated evolution arising from linkage or selection from spurious correlations arising due to chance.

In an effort to fill this gap, we have extended a computational approach, implemented in a software called AEGIS (Analysis of Epistasis and Genomic Interacting Sites), designed to detect both positively and negatively correlated pairs of mutations with very high specificity (Nshogozabahizi *et al.*, 2017). Although the original approach focused on identifying correlated evolution among sites within genes (Ibeh *et al.*, 2016; Nshogozabahizi *et al.*, 2017; Aris-Brosou *et al.*, 2017), nothing prevents it from being used to perform the same task across entire genomes. AEGIS makes use of Pagel’s phylogenetically informed maximum likelihood model for predicting correlated evolution between pairs of traits based on the co-distribution of their values (Pagel, 1994), and was extensively tested through both simulations and analyses of real data in the context of detecting correlated evolution at the molecular level, between pairs of nucleotide positions in a particular genome alignment (Nshogozabahizi *et al.*, 2017). The output of AEGIS is a list of pairs of sites and their associated probabilities of co-evolving by chance. While this approach uses phylogenetic information to reduce the detection of spurious associations, it is agnostic to the underlying mechanism responsible for correlation among sites, and thus cannot distinguish between linkage and selection leading to nonadditive fitness effects, or epistasis, as the cause of correlated evolution. This approach, like most statistical comparative methods, are now known to detect correlated evolution even in the case of a single unreplicated co-distribution of traits (Maddison and FitzJohn, 2014; Uyeda *et al.*, 2018).

Consequently, identifying the mechanisms driving correlated evolution still requires direct experimental measurements of epistasis between sites, preferably under conditions similar to the selective setting in which the pairs of sites evolved. To do this, we focused on the evolution of resistance to fluoroquinolone antibiotics in the opportunistic pathogen *Pseudomonas aeruginosa*. This pathogen is a ubiquitous, Gram-negative bacterium that causes both acute opportunistic infections and chronic respiratory tract infections in cystic fibrosis patients. Treatment with fluoroquinolone antibiotics imposes strong selection that regularly leads to the evolution of resistance at a number of well-known target sites such as efflux pump regulators (e.g. *nfxB*) and DNA topoisomerases (i.e. *gyrA*, *parC*) (Akasaka *et al.*, 2001; Wong *et al.*, 2012; Melnyk *et al.*, 2015; Kos *et al.*, 2015). As *P. aeruginosa* is amenable to genetic manipulation, we can introduce putative correlated mutations, either one at a time or in combination, into a range of genetic backgrounds. The level of epistasis can then be measured by comparing the effects of the two substitutions against the expected combined effects of each single substitution on traits relevant to selection, such as antibiotic resistance or fitness in the absence of antibiotics.

Here, we use a unique data set of 393 *P. aeruginosa* exomes to predict correlated evolution among sites and evaluate the mechanistic causes of correlations that evolve during selection by fluoroquinolone antibiotics. We investigate the role of epistasis and linkage, both physical and through hitchhiking, as causes of correlated evolution using a combination of tools including site-directed mutagenesis to reconstruct single and double mutants. For nonsynonymous substitutions that co-evolved in response to antibiotic selection, we test for epistasis with direct estimates of the minimum inhibitory concentration (MIC) of antibiotics and competitive fitness in the absence of drugs. We employ additional computational analyses to infer the mechanism leading to the correlated evolution of synonymous substitutions. Our results demonstrate that epistasis can drive correlated evolution among nonsynonymous sites tied to antibiotic resistance, whereas correlated evolution among synonymous sites is best explained by idiosyncratic processes.

## 2 MATERIALS AND METHODS

### 2.1 **Alignment and phylogeny**

We assembled a multiple sequence alignment which included the complete genomes of *P. aeruginosa* strains PA01 (PA01), *P. aeruginosa* UCBPP PA14 (PA14), and *P. aeruginosa* PA7 (PA7) downloaded from www.pseudomonas.com (Winsor *et al.*, 2016, accessed Dec 4, 2014), as well as 390 draft *P. aeruginosa* genomes (Kos *et al.*, 2015), used here as they were published alongside information detailing which of the strains were resistant to the fluoroquinolone antibiotic levofloxacin. The genomes used in our study represent a collection of clinical strains isolated from around the world (Table S1). Due to dissimilarities among contigs across the draft genomes, we were unable to construct a global alignment using whole genome alignment algorithms as implemented in either MAUVE (Darling *et al.*, 2004) or MUGSY (Angiuoli and Salzberg, 2011). Using PA14 reference gene sequences detailed in a database downloaded from www.pseudomonas.com (Winsor *et al.*, 2016, accessed Dec 4, 2014), we assembled an exome alignment using a custom algorithm to concatenate individual gene alignments.

**Table 1.** Results from MIC assays of mutant constructs under ciprofloxacin. We measured MIC from four technical replicates for each of two independent constructs (biological replicates), except for *gyrA c*248*t* that had three independent constructs. The significance of differences in log_2_ MIC between wild-type and each mutant construct was assessed with the Dunn test and a Bonferonni correction; *P* 0.05 shown in bold. Epistasis was measured with a multiplicative model, using the MIC determined from reads of the optical density (600 nm) after 24 hours growth, and measurement error was calculated using error propagation (Trindade *et al.*, 2009). There is evidence for epistasis when the absolute value of *e* is greater than the error of our measures (in bold).

| Background | Mutant | $X^2$ | Z | P | $\epsilon$ | Error |
| --- | --- | --- | --- | --- | --- | --- |
| PA14 | *gyrA c248t* | 6.480 | -2.546 | **0.005** |  |  |
|  | *parC c260t* | 0.000 | 0.000 | 0.500 |  |  |
|  | *parC c260g* | 0.000 | 0.000 | 0.500 |  |  |
|  | *gyrA c248t parC c260t* | 5.478 | -2.341 | **0.010** | **0.547** | 0.382 |
|  | *gyrA c248t parC c260g* | 5.478 | -2.341 | **0.010** | **0.547** | 0.382 |
| PA01 | *gyrA c248t* | 4.253 | -2.062 | 0.020 |  |  |
|  | *parC c260t* | 0.000 | 0.000 | 0.500 |  |  |
|  | *parC c260g* | 0.000 | 0.000 | 0.500 |  |  |
|  | *gyrA c248t parC c260t* | 4.000 | -2.000 | **0.023** | **4.461** | 2.051 |
|  | *gyrA c248t parC c260g* | 3.529 | -1.879 | **0.030** | **3.271** | 3.138 |

The species tree of these bacterial genomes was reconstructed based on ‘highly networked’ genes that, according to the complexity hypothesis, are unlikely to be horizontally transferred. We identified these highly networked genes from the Cluster of Orthologous Groups database’s (Tatusov *et al.*, 1997; Galperin *et al.*, 2015) information class genes (those with COG terms A, B, J, K, and L). From an alignment of 1,290 highly networked *P. aeruginosa* genes, we estimated the species tree by maximum likelihood with FastTree (Price *et al.*, 2010) (GTR + r model of evolution (Aris-Brosou and Rodrigue, 2012)) compiled with *DUSE_DOUBLE* as recommended for large sequence alignments. This tree was rooted with the subclade containing the taxonomic outliers PA7 (Roy *et al.*, 2010), AZPAE14941, AZPAE14901 and AZPAE15042 (Kos *et al.*, 2015). After rooting the tree, we removed the subclade from both the phylogeny and the above exome alignment, so that downstream analyses were based on 389 genomes. For additional information please see *Alignment algorithm* and *Complexity Hypothesis* in the supplementary information.

### 2.2 **Analysis of correlated evolution**

We used AEGIS (Nshogozabahizi *et al.*, 2017) to detect pairs of nucleotide positions (sites) that evolved in a correlated manner. For this, AEGIS performed pairwise comparisons between sites, testing if a model of dependent evolution was significantly better than an independent model at explaining the observed phylogenetic distribution of nucleotide states. This maximum likelihood analysis relied on the algorithm (Pagel, 1994), computationally validated with AEGIS for use with amino acid data (Nshogozabahizi *et al.*, 2017), as implemented in BayesTraits for binary states (Pagel and Meade, 2006). Nucleotide states in our alignment were converted into binary characters where ‘0’ was assigned where there was a consensus of nucleotide character at 80% of the levofloxacin sensitive strains and ‘1’ otherwise. We chose this method of recoding so that signals of correlated evolution would more likely reflect adaptation to fluoroquinolone antibiotic selection. All R scripts and complete results can be found at github.com/JDench/sigResults_AEGIS_inVivo. Due to computational limitations, we limited our analysis of correlated evolution to a 12 focal gene by exome analysis. Six of the focal genes were previously shown to evolve during fluoroquinolone selection (Akasaka *et al.*, 2001; Wong *et al.*, 2012) (*gyrA*, *gyrB*, *nfxB*, *parC*, *parE*) or are linked to biofilm formation (Choy *et al.*, 2004), which can contribute to antibiotic resistance (Starkey *et al.*, 2009) (*morA*). The six other genes (*dnaA*, *dnaN*, *lpd3*, *ribD*, *rpoB*, *serC*) were randomly selected from among the other 5,971 genes in our alignment. For additional information please see *Analysis of correlated evolution* in the supplementary information.

### 2.3 Direct experimental assessment of epistasis

All growth and incubation conditions were performed in lysogeny broth (LB) (Bertani, 1951) containing half the normal salt (i.e., 5g/L NaCl), at 37°C, in an orbital shaker set at 150 rpm.

**Generating mutants:** To measure the *in vitro* fitness and antibiotic resistance effects of substitutions predicted with AEGIS, we created mutant constructs using a modified selection counter-selection allelic replacement protocol (Melnyk *et al.*, 2017). Here, replacement alleles were constructed from the wild-type (WT) template and contained only the mutations of interest rather than coming from an evolved strain which may carry non-focal mutations. As the effect of a mutation may be contingent upon the genetic background (i.e., WT) in which it arose (Sorrells *et al.*, 2015; Kryazhimskiy *et al.*, 2014; Blount *et al.*, 2008), constructs were created using two distinct genetic backgrounds: PA01, and PA14 (Dettman *et al.*, 2013). For additional information please see *Modified selection counter-selection protocol* in the supplementary information.

**Minimum inhibitory concentration assays:** We performed minimum inhibitory concentration assays to identify if our mutant constructs had increased resistance to the fluoroquinolone antibiotic ciprofloxacin. Using a 96-well plate, we carried out serial 2-fold dilutions of ciprofloxacin, in a liquid LB broth, such that columns represented a concentration gradient from 32 to 0.015625 (*i.e.* 2^(5,4,…,−6)^) *µ*g/mL. We inoculated each well with 5 *µ*L of overnight mutant culture such that each row represented a single type of inoculum. After 24 hours of growth, we read the absorbance values (at 600 nm) using a plate reader (Table S7). Each plate contained at least one blank row, used as reference for reads of the absorbance. We performed four replicates of each genotype, while ensuring that no genotype appeared on a single plate more than once.

**Competitive fitness assays:** We measured relative fitness of WT and mutant constructs via competitive fitness assays. Competitions began when we mixed an equal volume of overnight culture, diluted 100 fold, of a focal strain and a marked competitor (Lenski *et al.*, 1991). Similar to previous work, the marked competitor was a WT strain bearing the neutral *lacZ* marker gene. The frequency of focal and marked strains were estimated using direct counts of diluted culture plated immediately after mixing and again after competition during overnight growth to stationary phase. We plated six replicates of each competition, and time point, on minimal media agar plates containing X-Gal. Counting was done after overnight growth at 37°C, followed by 2-4 days of growth at room temperature. For additional information please see *Calculating competitive fitness* in the supplementary information.

### 2.4 Quantifying importance of sites in predicting resistance

We quantified the importance of polymorphic sites in our alignment for predicting resistance to levofloxacin based on the machine learning classification algorithm ‘adaptive boosting’, implemented in the *adaBoost* (Alfaro *et al.*, 2013) R package. Measurements of importance provided a relative measure of the strength of association between resistance and substitution at a site in our alignment. We compared the machine learning results to substitutions known to be within the six, previously identified, focal *P. aeruginosa* genes of which five evolve in response to fluoroquinolone selection (Akasaka *et al.*, 2001; Wong *et al.*, 2012; Kos *et al.*, 2015) and one is involved in biofilm formation (Choy *et al.*, 2004) which can contribute to antibiotic resistance (Starkey *et al.*, 2009). For additional information please see *adaptive boosting algorithm* in the supplementary information.

### 2.5 Assessing horizontal gene transfer

Horizontal gene transfer (HGT) was assessed with a phylogenetic approach (Ravenhall *et al.*, 2015), where we reconstructed each gene tree as above (maximum likelihood under GTR + r), and tested for tree concordance with the Approximately Unbiased (AU) test (Shimodaira, 2002). This was done with CONSEL (Shimodaira and Hasegawa, 2001), on the *baseml* (GTR + r_5_, no clock) output from PAML (Yang, 2007).

### 2.6 Testing for biological effects of synonymous substitutions

To test if pairs of synonymous substitutions were correlated due to translational effects, we estimated the free energy (Δ*G*) of folded mRNA transcripts for both WT and mutant transcripts, which we normalized as relative values through division of the WT estimate (Δ*G_rel_*). We also estimated if synonymous mutations affected translation elongation rates via estimates of the index of translation elongation (*I_T E_*: Xia, 2014). For additional information please see *Calculating* Δ*G and I_T E_* in the supplementary information.

## 3 RESULTS AND DISCUSSION

### 3.1 **Predicting correlated evolution among sites**

We identified pairs of sites that evolve in a correlated manner in the context of fluoroquinolone resistance by first assembling a multiple sequence alignment using the entire set of 5,977 proteincoding genes from 393 previously sequenced *P. aeruginosa* strains. This multiple sequence alignment included three reference strains (PA01, PA14, and PA7) and 390 clinical strains for which we had information on the level of levofloxacin (a fluoroquinolone antibiotic) resistance (Kos *et al.*, 2015). We then used this alignment to construct a phylogenetic tree rooted by PA7, an outlier strain (Roy *et al.*, 2010) (Fig. S2). The subclade containing this strain was removed for downstream analyses. We acknowledge that the comparative approach implemented in AEGIS may not be ideally suited to analyze highly promiscuous bacteria; however, we constructed our phylogeny from genes unlikely to undergo horizontal transfer so that signals of correlated evolution would represent independent evolutionary events.

We subsequently employed AEGIS to identify pairs of nucleotide positions that show evidence of correlated evolution. As this approach can be computationally expensive, requiring on the order of *n*^2^ comparisons in a genome of length *n* (“’ 84 × 10^9^ comparisons), we focused our analyses on a subset of twelve genes against all other positions in the exome. AEGIS calculated the probability that each position within these twelve genes evolved in a correlated manner with any other nucleotide position in the entire *P. aeruginosa* exome. Among the more than 144 million pairwise comparisons, we found that “’ 127, 000 pairs of sites showed evidence of significant correlation with *P* ≤ 0.01 after a Benjamini-Hochberg correction for false discovery rate (FDR). At medium (*P* ≤ 10^−7^) or high (*P* ≤ 10^−11^) significance levels, as previously determined based on extensive simulations (Nshogozabahizi *et al.*, 2017), only 178 and nine pairs, respectively, showed evidence for correlated evolution (Fig. 1). Among the nine pairs of sites showing the strongest evidence for correlated evolution, we found substitutions in the nucleotide sequence for each of the two DNA gyrases (*gyrA* and *gyrB*) targeted by fluoroquinolones, within the canonical fluoroquinolone resistance determining gene *parC*, in the biofilm-associated gene *morA*, but also in *dnaN* which was among the set of randomly chosen genes. Notably, all highly significant pairs of substitutions were synonymous, and hence unlikely to affect drug resistance or fitness (but see Plotkin and Kudla, 2011; Bailey *et al.*, 2014; Agashe *et al.*, 2016), save for one pair of nonsynonymous sites, *gyrA* (c248t)-*parC* (c260g/t), that presumably impacts the level of resistance (Fig. 1). The next most strongly correlated pair of nonsynonymous substitutions involved *parC* and an uncharacterized gene at locus tag *PA14_34000*, but its significance was several orders of magnitude lower than any of the nine strongest correlated pairs (*P* = 8.018 × 10^−7^ here *vs. P* ≤ 10^−11^ above). In total, we found evidence of correlated evolution among 96 paired nonsynonymous substitutions with *P* ≤ 10^−4^, and many more with *P* ≤ 10^−2^. However, most involved uncharacterized genes while the rest involved genes without sufficient biological rationale to justify investigation within our study. It should be noted that the algorithm we employed to detect correlated evolution (*i.e.* Pagel’s algorithm Pagel, 1994), just like pretty much any alternative phylogeny-aware methods used in comparative studies (*e.g.*, Maddison, 1990; Huelsenbeck *et al.*, 2003; Maddison *et al.*, 2007), can detect a significant statistical association even when there is a single case of co-distribution of traits (Maddison and FitzJohn, 2014; Uyeda *et al.*, 2018). This is why, despite the care taken in our approach, we performed additional experimental analyses to validate the statistical signal detected between the most strongly correlated pairs.

**Figure 1.**
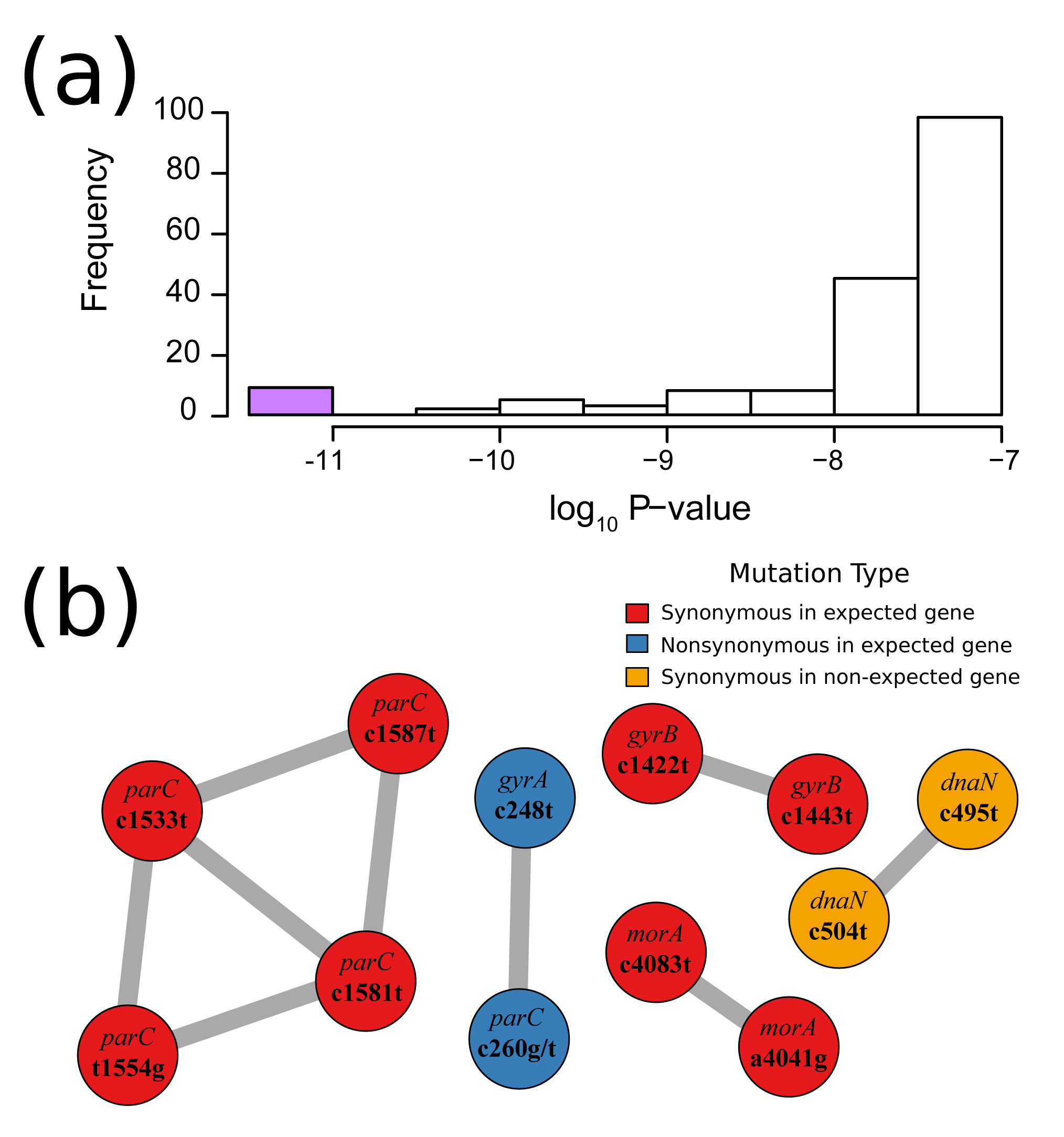
Correlated pairs of mutations with strongest evidence. (a) a histogram showing the distribution of significance for all correlated pairs with strong (*P <* 10^−7^) evidence for correlated evolution. The purple bar highlights the strength of evidence for the pairs presented in B. (b) substitutions with the strongest evidence for correlated evolution (*P <* 10^−11^). Circles represent a substitution in a gene (top text), at a particular nucleotide position (bottom text) on the coding strand. Colours show whether or not the substitution was synonymous, and whether or not it was found in one of the six genes expected to evolve in response to fluoroquinolone selection. Edges connect the predicted pairs of correlated substitutions.

### 3.2 Experimental validation of the nonsynonymous pair

To test if epistasis was driving the correlated evolution of these fluoroquinolone resistance substitutions in *gyrA-parC*, we used a modified allelic replacement protocol (Melnyk *et al.*, 2017) to construct mutants bearing all possible combinations of *gyrA-parC* single and double mutations in two genetic backgrounds (PA01 and PA14) that lack the mutations of interest (Methods). We then quantified the MIC of a fluoroquinolone antibiotic (ciprofloxacin) for all genotypes and their competitive fitness against the ancestral (wild-type [WT]) genotypes in the absence of drug. We used the PA01 and PA14 genetic backgrounds because they are phylogenetically distinct (Dettman *et al.*, 2013), representing two major clades (Kos *et al.*, 2015).

Our results reveal that the pattern of changes in resistance for single and double mutants was similar for both genotypes (Fig. 2a,b). On its own, the *gyrA* mutation increases resistance, by “’ 8-fold in PA14 and at least twice that in PA01, whereas the *parC* mutations on their own have no effect. In combination, however, the *gyrA* mutation together with either of the *parC* mutations confers between 128-256 fold increases in resistance, depending on genetic background. Support for this interpretation comes from a likelihood ratio test for significant fixed effects in a mixed effects model using biological replicates as a random effect. Results of the tests show that MIC depends on both the genetic background (*X*^2^= 46.52, *df* = 1, *P* = 9.059 ×10^−12^) and mutation (*X*^2^ = 813.95, *df* = 6, *P <* 2.2 × 10^−16^), but also that the magnitude of the fold-increase in MIC was similar for PA01 and PA14 constructs (i.e. interaction term, *X*^2^ = 7.47, *df* = 5, *P* = 0.1878). This analysis confirms the presence of strong positive epistasis for resistance between these *gyrA-parC* mutants, the magnitude of which is independent of the two genetic backgrounds tested.

**Figure 2.**
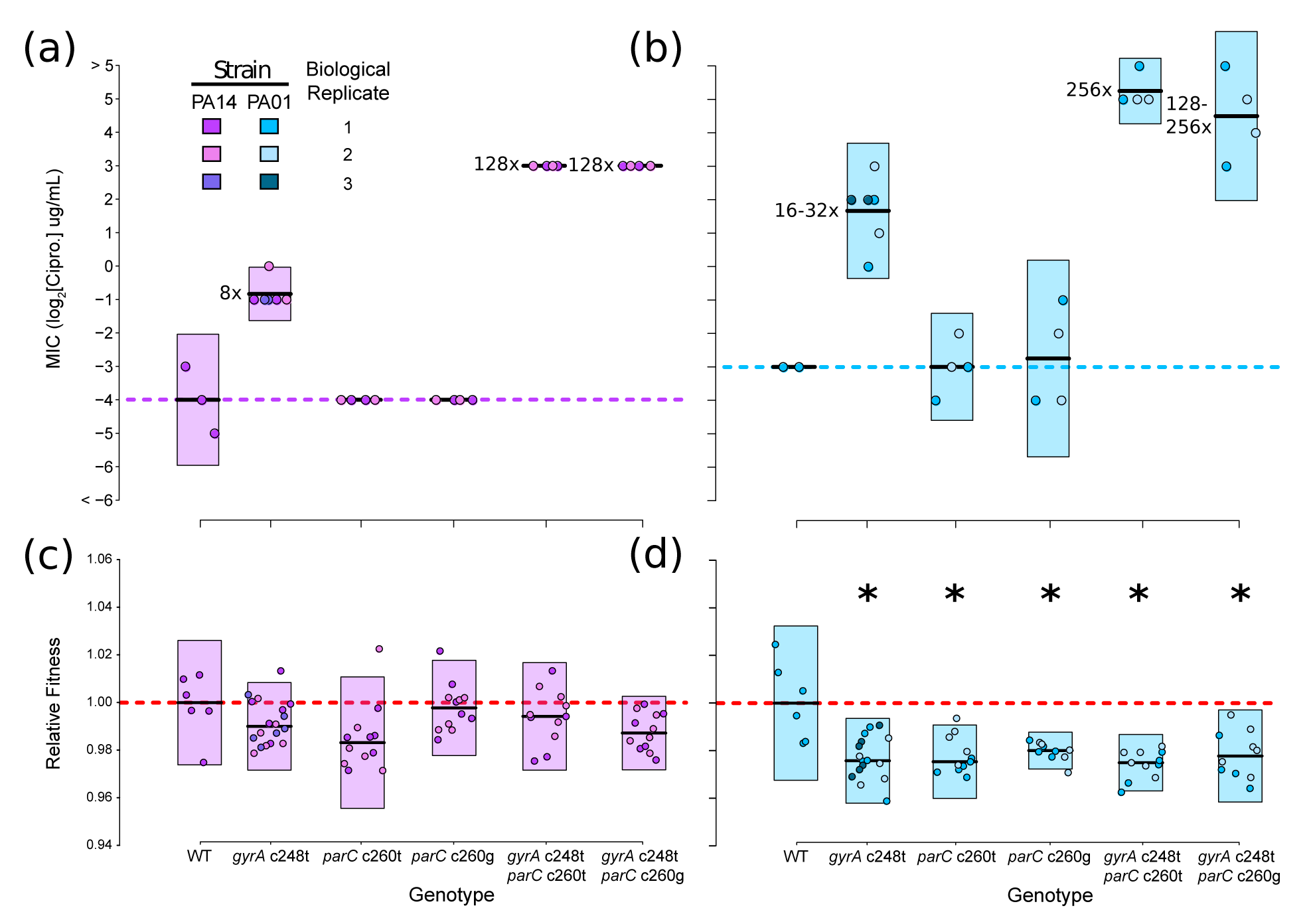
Empirical tests of epistasis for resistance and fitness. The mean and 95% confidence interval of measurements for each genotype is represented by the dark black lines and colored rectangles respectively. The color used reflects the genetic background of genotypes while the hue identifies the biological replicate. (a,b) results of the ciprofloxacin MIC assay of *P. aeruginosa* WT and mutant constructs. Dashed horizontal lines highlight the MIC value for WT strains. Numbers to the left of mean (black) lines represent the MIC fold increase, compared to WT, of mutants. (c,d) results of the competitive fitness assays for WT and mutant construct genotypes. The relative fitness of WT strains is highlighted by the horizontal dashed red line. Mutants with relative fitness significantly different from wild-type are denoted with an asterisk (see Table 2).

**Table 2.**
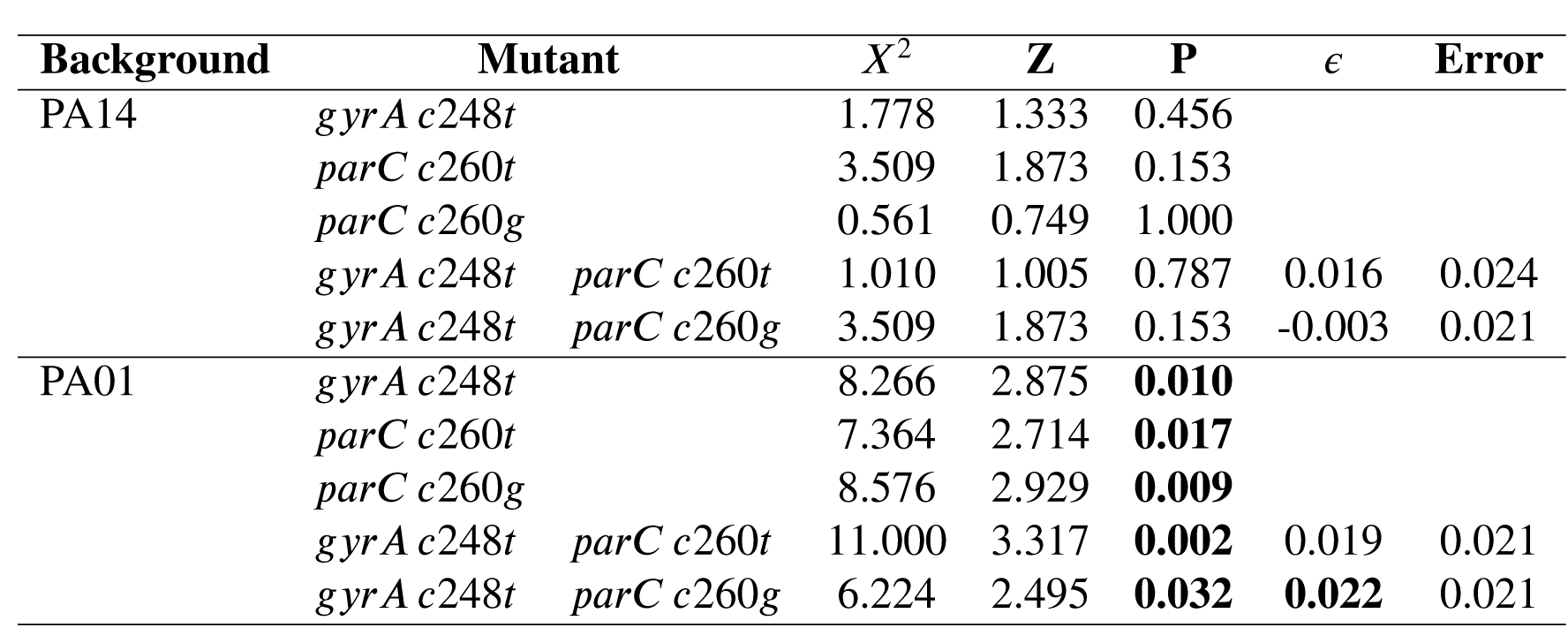
Results from competitive fitness assays of mutant constructs in permissive LB media. We measured fitness from six technical replicates for each of two independent constructs (biological replicates), except for *gyrA c*248*t* which had three independent constructs. Relative fitness was calculated by dividing the fitness of each mutant construct by that of its wild-type (not shown here). The significance of differences in competitive fitness of wild-type and each mutant construct was assessed with the Dunn test and a Bonferonni correction; *P* 0.05 shown in bold. Epistasis was measured with a multiplicative model and error was calculated using error propagation (Trindade *et al.*, 2009). There is evidence for epistasis when the absolute value of *e* is greater than the error of our measures (in bold).

| Background | Mutant | $X^2$ | Z | P | $\epsilon$ | Error |
| --- | --- | --- | --- | --- | --- | --- |
| PA14 | *gyrA c248t* | 1.778 | 1.333 | 0.456 |  |  |
|  | *parC c260t* | 3.509 | 1.873 | 0.153 |  |  |
|  | *parC c260g* | 0.561 | 0.749 | 1.000 |  |  |
|  | *gyrA c248t parC c260t* | 1.010 | 1.005 | 0.787 | 0.016 | 0.024 |
|  | *gyrA c248t parC c260g* | 3.509 | 1.873 | 0.153 | -0.003 | 0.021 |
| PA01 | *gyrA c248t* | 8.266 | 2.875 | **0.010** |  |  |
|  | *parC c260t* | 7.364 | 2.714 | **0.017** |  |  |
|  | *parC c260g* | 8.576 | 2.929 | **0.009** |  |  |
|  | *gyrA c248t parC c260t* | 11.000 | 3.317 | **0.002** | 0.019 | 0.021 |
|  | *gyrA c248t parC c260g* | 6.224 | 2.495 | **0.032** | **0.022** | 0.021 |

Notably, mutants carrying the *gyrA c248t* substitution had the expected 8-32 fold increase in ciprofloxacin resistance (Akasaka *et al.*, 2001). However, the *parC c260g/t* substitutions did not affect resistance on their own but did show a “’ 16 fold additional increase for double mutants (Fig. 2 a,b; Table 1). This strong epistasis for resistance could explain why *gyrA c248t* and *parC c260g/t* double mutants are common among fluoroquinolone resistant clinical isolates of *P. aeruginosa* (Akasaka *et al.*, 2001; Kos *et al.*, 2015) and is consistent with previous work showing that the fluoroquinolone resistance conferred by substitutions in *parC* depended upon mutations in *gyrA* (Moon *et al.*, 2010). We note that Akasaka *et al.* (2001) quantified the effect on MIC of both the *gyrA* and *parC* mutations by measuring decatenation activity of purified protein and found that both mutations would increase resistance. This result suggests a possible Bateson-like molecular mechanism of the observed epistasis for resistance (Bateson, 1911). As ciprofloxacin acts by inhibiting both DNA gyrase and topoisomerase IV (the protein products of *gyrA* and *parC*, respectively), the changed decatenation activity of topoisomerase IV caused by the *parC* substitution may be masked if DNA replication is prevented by non-functional DNA gyrase. Since *gyrA* single mutants can grow under elevated fluoroquinolones levels, but *parC* single mutants cannot, it is reasonable to posit that the *gyrA* mutation compensates for the inhibition of *parC*, or that fluoroquinolones do not entirely inhibit *parC* protein activity. A study of the molecular basis for this epistasis for resistance may provide valuable insight in the development of next generation fluoroquinolones.

The results for fitness costs associated with resistance mutations tell quite a different story (Fig. 2c,d). We found no evidence for a cost of resistance associated with the *gyrA* and *parC* mu tations, either singly or in combination, in the PA14 background, a result that has been observed previously (Melnyk *et al.*, 2015). By contrast, all the single and double mutations in PA01 lead to significant fitness costs, and the double mutant including the *parC c260g* mutation, exhibits significant positive epistasis (Table 2). In other words, the resistance mutations compensate for each other, leading to costs that are lower than expected from their additive effects. Our results support the idea that the cost of antibiotic resistance in bacteria can vary substantially across genetic backgrounds (Melnyk *et al.*, 2015). Combinations of mutations in *gyrA* and *parC* other than those tested here have been shown to be cost-free in *Streptococcus pneumoniae* (Gillespie *et al.*, 2002), reinforcing the idea that the effect of a mutation, or combination of mutations, depends intimately on genetic background. Moreover, our observation of positive epistasis for resistance in both genetic backgrounds, but not consistently for fitness costs, indicates that the correlated evolution detected in *gyrA* and *parC* is likely driven by epistatic effects on antibiotic resistance, rather than competitive fitness in antibiotic-free environments.

Six of the genes in our twelve gene-by-exome analysis were expected to evolve in response to fluoroquinolone selection. Yet, we only had sufficient evidence to support experimental validation of epistasis between a single pair of sites canonically associated with fluoroquinolone resistance mutations. Two questions remained: (i) how come AEGIS did not identify correlated evolution involving additional nonsynonymous resistance mutations, and (ii) what was/were the mechanism/-s underlying the highly significant correlation identified for the eight pairs of synonymous substitutions?

### 3.3 Resistance is strongly associated with only two substitutions in our alignment

The computational approach employed above identified only two nonsynonymous substitutions likely to be correlated during the evolution of resistance to fluoroquinolone antibiotics (Wi- bowo, 2013; Kos *et al.*, 2015). Our lack of success in identifying more pairs of nonsynonymous substitutions could be because no other resistance mutations in our alignment evolved in a correlated manner, or because AEGIS is only able to detect instances of strong correlated evolution (Nshogozabahizi *et al.*, 2017), or perhaps that only a few substitutions in our dataset are strongly associated with drug resistance. We therefore performed three additional analyses to uncover further substitutions associated with drug resistance in our dataset.

First, we quantified the importance of all polymorphic positions in the *P. aeruginosa* exome for predicting fluoroquinolone resistance (Long *et al.*, 2019). We did this by training a supervised machine-learning algorithm, based on adaptive boosting, which measured the strength of correlation between substitutions in our alignment and the resistance phenotype. This approach detected that the *gyrA 248* and *parC 260* nucleotide positions were the only two sites in our alignment that strongly predicted resistance (Fig. S4). These two positions were also the only ones among the 100 most important associations that were located in genes expected to evolve in response to fluoroquinolone selection (Fig. S4).

Second, we tested the association between fluoroquinolone resistance and nonsynonymous mutations at known fluoroquinolone resistance determining sites (Akasaka *et al.*, 2001; Kos *et al.*, 2015). From our alignment, the *gyrA 248* (*T83I*) and *parC 260* (*S87L* and *S87W*) mutations were present in 41.4% and 32.9% of all strains, respectively, and these mutations were correlated to the resistance phenotype (*χ*^2^ tests: *gyrA 248*, *X*^2^ = 231.941, *df* = 1, *P* = 2.250 × 10^−52^; *parC 260*, *X*^2^ = 177.21, *df* = 1, *P* = 1.98 × 10^−40^, Table S2). We note that another study has shown evidence for epistasis in the context of drug resistance when mutations in *parC* (*S87L* and *S87W*) co-occur with a mutation in *gyrA*, but at a different position (*S91F* instead of *T83I*; Schubert *et al.*, 2019). In contrast, nonsynonymous substitutions at additional amino acid sites known to confer fluoroquinolone resistance were identified in our alignment, but either did not correlate with resistance (at *P* ≤ 0.05), or represented less than 10% of all strains included in our analysis (max. 8.74%, mean (sd) 3.54 (1.81)%; Table S2). Previous work showed that the signal of correlated evolution (i.e. the sensitivity of the AEGIS algorithm) is maximized when roughly half the strains are mutants, and when these mutations are evenly distributed throughout the phylogeny (Nshogozabahizi *et al.*, 2017), which is not the case here.

Third, we searched for loss of function mutations that are known to occur under fluoroquinolone selection, such as those affecting the *nfxB* or *orfN* genes (Wong *et al.*, 2012). There were no examples of premature stop codons within the sequences of these two genes in our alignment, although the presence of nonsynonymous loss of function mutations cannot be ruled out. Although it is in principle possible to resort to branch-site codon models to identify positions under selection in branches of interest (Yang and Nielsen, 2002; Zhang *et al.*, 2005), and hence detect sites potentially involved in drug resistance, our alignment contains too few taxa and too little evolutionary depth for proper statistical analysis (Arenas, 2015). Altogether, these three lines of evidence suggest that among the subset of known fluoroquinolone resistance mutations, only the *gyrA c248t* and *parC c260g/t* substitutions are present in enough strains to have been detected as correlated and show a significant association with fluoroquinolone resistance in our alignment.

### 3.4 Synonymous substitutions: the role of multiple drivers

AEGIS provided strong evidence for correlated evolution among synonymous mutations. This result is surprising because synonymous mutations are often assumed to evolve neutrally, and so we would not *a priori* expect them to exhibit strong correlated evolution unless they were under strong selection or were physically linked. This result could be due to strong phylogenetic uncertainty, which can affect comparative methods (Shoji *et al.*, 2016; Guigueno *et al.*, 2019). To account for this we could rerun the analyses on bootstrapped trees (Nshogozabahizi *et al.*, 2017) but this is a taxing solution as it would be computationally quite demanding, especially here, possessing statistical properties that would deserve scrutiny at a level beyond the scope of the present work. However, the phylogenetic tree that we reconstructed has internal nodes that are fairly well supported (Fig. S2), making it unlikely that the signal of correlated evolution that we detect between synonymous mutations be due to phylogenetic uncertainty.

On the other hand, it has been known for some time that synonymous sites can experience purifying selection (Zhou *et al.*, 2010; Lawrie *et al.*, 2013), and a number of recent reports show that they can sometimes contribute to adaptation as well (Bailey *et al.*, 2014; Agashe *et al.*, 2016). However, we find no signal of epistasis-driven selection acting on the most strongly correlated pairs of synonymous sites identified in our study (Fig. 1). Estimates of the change in stability of mRNA transcripts and index of translation efficiency, which are proxies for translation rate (Kudla *et al.*, 2009) and efficiency (Xia, 2014), respectively, relative to the wild type are shown in Fig. 3 and Tables S2,3. While two of the synonymous mutations in *parC* (*c*1581*t* and *c*1587*t*) could produce more stable and abundant mRNA transcripts, the only evidence for epistasis in either metric we could identify was for translation efficiency in the mutations in *morA*, and the direction of this effect is opposite to what would be expected if it were to result in positive selection.

**Figure 3.**
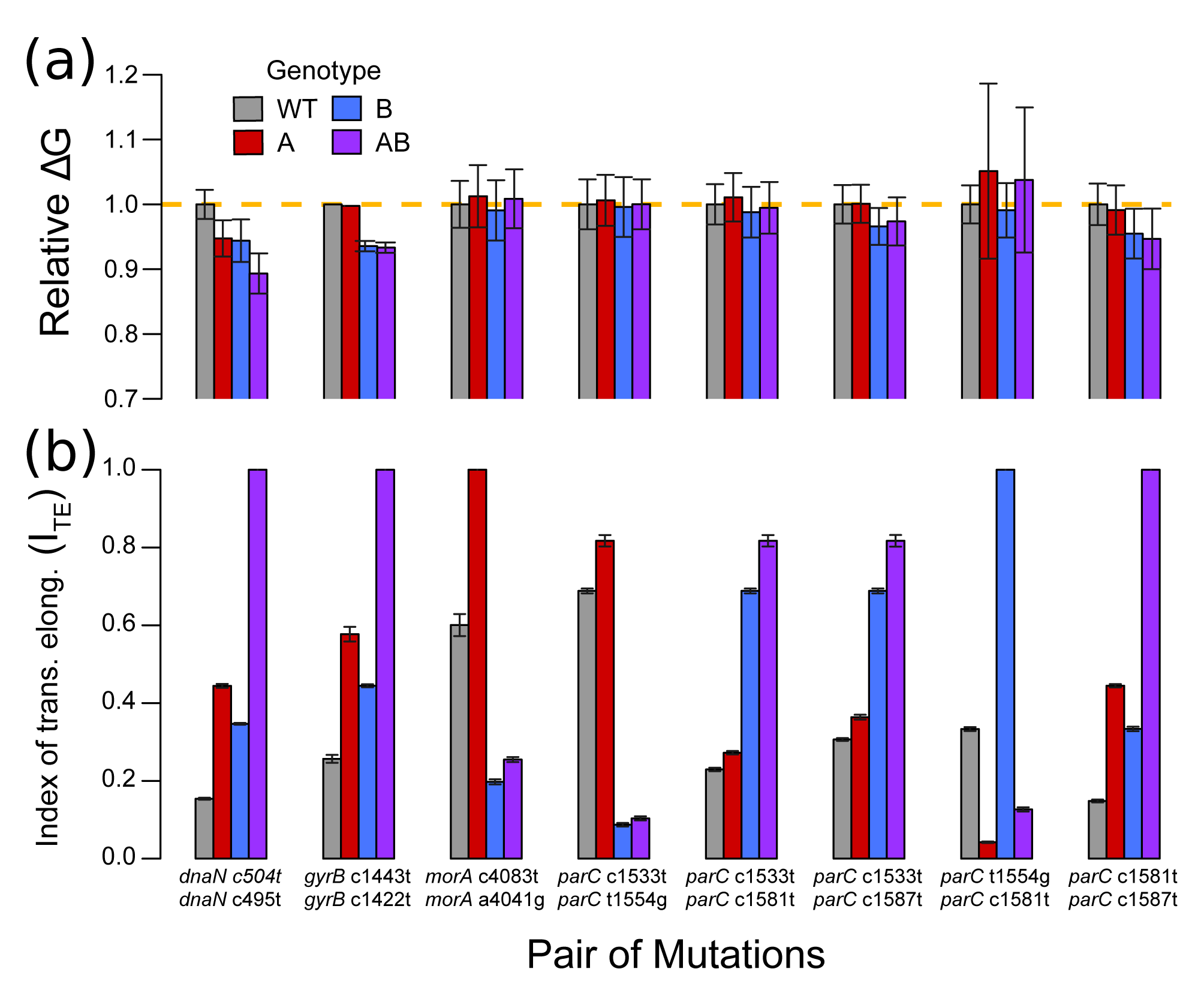
Computationally predicted biological effects of the strongly correlated pairs of synonymous substitutions. Each set of bars is for a pair of substitutions, where the first bar (grey) is the wild-type value (inferred from PA14 reference genome), the second bar (red) is a mutant carrying the first substitution (A) listed below the bars while the third bar (blue) represents a mutant carrying the second listed substitution (B), and the fourth bar (purple) represents a double mutant. (a) estimates of the relative change of mRNA folding free energy (Δ*G*) are relative to the WT value (gold dashed line) and error bars represent the 95% confidence interval. (b) measures of the mean index of translation elongation (I_TE_, B) with error bars presenting standard deviation among measures estimated from double mutant strains present in our data set.

**Table 3.**
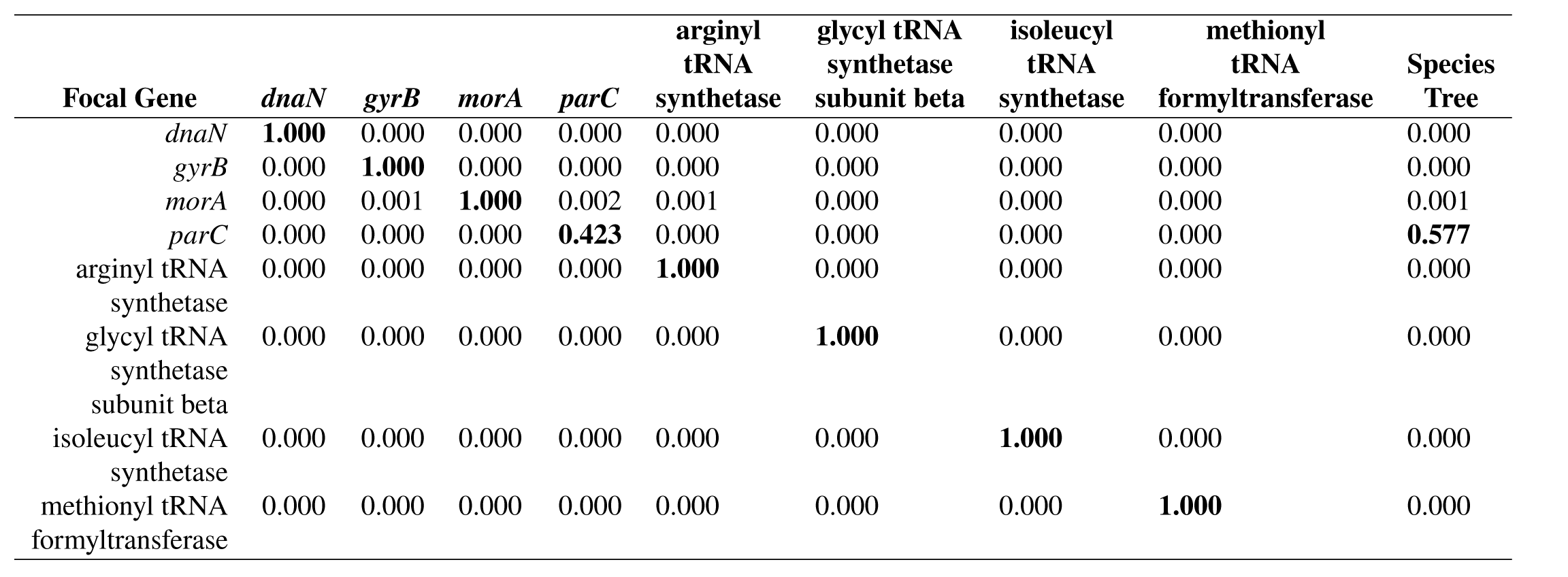
Approximately unbiased (AU) tests of significant differences among phylogenetic trees. We built each tree using the multiple sequence alignment for one of the four focal genes that contain the most significantly correlated synonymous substitutions (*dnaN*, *gyrB*, *morA*, *parC*) and four tRNA genes unlikely, according to the complexity hypothesis (Jain *et al.*, 1999; Aris-Brosou, 2005) to undergo HGT (*arginyl tRNA synthetase, glycyl tRNA synthetase subunit beta, isoleucyl tRNA synthetase, methionyl tRNA formyltransferase*). We tested for significant differences among all the gene trees and the species tree used in the analysis of correlated evolution. Values in each cell represent the probability, calculated via the AU test, that a tree (column names) describes evolution observed in an alignment (row names). In bold are the “best” tree(s) (*i.e.:* insignificantly different trees).

By contrast, two lines of evidence point to physical linkage between sites as the proximate cause of correlation between synonymous mutations. First, the proportion of synonymous pairs occurring within the same gene was higher than expected by chance among all observed (synonymous and nonsynonymous) pairs (*X*^2^ = 7246.037, *df* = 2, *P* ≤ 2.16 × 10^−16^; see Table S6). Second, the physical distance between pairs of correlated synonymous substitutions, as mapped on the circular genome of PA14, was negatively associated with the strength of correlation (for pairs with *P* ≤ 10^−4^; *t* = −5.1784, *df* = 7, 528, *P* = 2.29 × 10^−7^ Fig. S5), though the correlation was quite poor (*R*^2^ = 0.03). Physical distance remains significantly associated to strength of correlation even when the 8 most significantly correlated pairs are removed substitutions (*t* = −6.922, *df* = 7, 520, *P* ≈ 4.8 × 10^−12^, *R*^2^ = 0.006).

The mechanism responsible for the close physical linkage between synonymous mutations is less obvious. One possibility is that both hitchhiked to high frequency alongside a third, non-synonymous mutation in the same gene that was under strong positive selection. We found a non-synonymous mutation at nucleotide position 1374 of *gyrB*, which is near the gene’s quinolone-resistance determining region, that was always present alongside the synonymous mutations. Moreover 97% of the strains that have substituted the pair of synonymous mutations in *morA* also fixed nonsynonymous mutations at other positions along the gene that could have functional effects on regulating or sensing internal oxygen levels (positions 292, 293, 306) (Taylor and Zhulin, 1999) or signal transduction (position 1356) (The UniProt Consortium, 2017). While these results point to hitchhiking as a plausible explanation for the correlation between sites, we failed to detect additional functional links with nonsynonymous mutations in the other genes. Moreover, AEGIS did not detect correlations between any of the correlated synonymous mutations and additional nonsynonymous mutations, even at *P* ≤ 10^−4^ (Fig. S1), suggesting that hitchhiking is not a general explanation for the correlated evolution of synonymous mutations in this data set. It should further be noted for the correlated pairs of substitutions found in *gyrB* and *morA*, that unless the pair of mutations repeatedly arose in the context of a third substitutions under selection, hitchhiking would not have produced a signal of correlated evolution.

A second possible mechanism we considered is horizontal gene transfer (HGT) and/or incomplete lineage sorting. Comparing the similarity of the species tree and the gene trees for each of the four genes with a strongly correlated pair of synonymous mutations reveals little correspondence between the two for all but *parC* (Table 3), suggesting widespread recombination in our strain collection. To further assess the prevalence of recombination, we included tRNA genes located in close proximity of each of our four focal genes in the PA14 assembly. The rationale for this analysis was that both tRNA genes and three of our four focal genes, being involved in information processing, should interact with a large number of other gene products (transcripts and/or proteins), so that a successful HGT of such genes should involve the co-transfer of most of their interacting partners, an unlikely event (Jain *et al.*, 1999; Aris-Brosou, 2005). However, we found that all tRNA gene trees differed from each other and the species tree (Table 3) suggesting that recombination (and thus HGT and/or incomplete lineage sorting) may be widespread in our strain collection. Widespread HGT is consistent with the findings of Kos *et al.* (2015), who found that the *parE* and *β*-lactamase genes were horizontally transferred in a number of the strains included in our exome alignment.

Importantly, both hitchhiking and HGT represent population-level processes that can lead to a pattern of physical linkage between synonymous sites. However, neither mechanism explains how such strong intra-gene correlations arose in the first place. Functional constraints, for example via intra-molecular pairing that forms strong hairpins in mRNA, and locally-biased mutation that depends on nucleotide context, have been proposed as potential mechanisms (Tsunoyama *et al.*, 2001). A full analysis of the contribution of these processes to the patterns we observe here, while interesting, is beyond the scope of this work, however.

Altogether, these results suggest that the close physical linkage between pairs of synonymous mutations is due in part to widespread HGT, except in the *parC* gene. Without any evidence that either HGT or hitchhiking with a nonsynonymous mutation are driving the correlated evolution of the four synonymous mutations in *parC*, which form a network of five correlated pairs (Fig. 1), these mutations could in principle have evolved repeatedly as a driverless cohort - that is, mutations that self-assemble by chance, and drift to fixation via linkage (Buskirk *et al.*, 2017). However, both the species tree (results of AEGIS analysis) and the gene trees (Fig. S3) show that these four synonymous *parC* substitutions have evolved independently numerous times. It is therefore unlikely that a cohort of four substitutions repeatedly self-assembled by chance. Whether these mutations co-occur for selective reasons related to the ecology of the CF lung (Sriramulu *et al.*, 2005; Smith *et al.*, 2006; Poole, 2005) and antibiotic treatment (Kos *et al.*, 2015) or other processes such as functional constraints associated with transcription and translation or mutational biases remains unclear.

## 4 CONCLUSIONS

Our results show many instances of correlated evolution (“’ 127, 000 interactions at *P* ≤ 0.01 from “’ 10, 220, 000 comparisons tested - *i.e.*: “’ 1.2%). In the context of antibiotic resistance in *P. aeruginosa*, moreover, some instances of correlated evolution can be underlain by, among other processes, epistasis. This result is consistent with experimental work showing that multiple mutations can contribute to the evolution of antibiotic resistance (Trindade *et al.*, 2009; MacLean *et al.*, 2010; Hall and MacLean, 2011), both within and between genes (Lehner, 2011), although typically the number of mutations identified remains quite small. Our results suggest, unsurprisingly perhaps, that a wide range of mutations can evolve in a correlated manner in the context of antibiotic selection – but also that only a limited number of these mutations are easily interpreted in the context of drug resistance (here, those found in *gyrA* and *parC*). Furthermore, our results lend support to the idea that high-throughput techniques for identifying correlated evolution *can* lead to identifying epistasis. Similar interpretations have been made previously, for instance in the case of the evolution of RNA genes where base pairing is critical (Dutheil *et al.*, 2010), or in the context of genome-wide association studies, where the detection of interacting residues is used as a first screen for epistasis, before fitting the linear models traditionally employed to identify genetic determinants of drug resistance (Schubert *et al.*, 2019). This latter approach, which notably also revealed epistatic interactions between *gyrA* and *parC*, requires that MIC values of all the genomes included in the analysis be known – data that was not available to us. In spite of this, both approaches should be highly complementary.

However correlated evolution is necessary but not sufficient evidence for epistasis (Nshogozabahizi *et al.*, 2017), as it can also arise through other mechanisms such as physical linkage. Computational approaches like ours detect relationships between phenotype and genomic substitutions though these associations are not evidence alone for a causal relationship. Indeed, we have observed a strong signal that physical linkage has driven the correlated evolution between pairs of synonymous substitutions in our data set, perhaps linked to the wholesale transfer of genes during recombination. Despite recent reports that synonymous mutations may not, in fact, be silent (Duan *et al.*, 2003; Chamary and Hurst, 2005; Plotkin and Kudla, 2011; Cuevas *et al.*, 2012; Bailey *et al.*, 2014; Agashe *et al.*, 2016; Fragata *et al.*, 2018), we have little evidence that synonymous substitutions, either on their own or in combination, impact fitness in our data. The fact that many of the most strongly correlated synonymous substitutions we identified here are GC---AT mutations (Fig. 3) reflects the pronounced GC bias of *P. aeruginosa* and reinforces the notion that these pairs of synonymous mutations have not been selected. It is notable that another computational study found evidence of physical linkage underlying correlated evolution between nonsynonymous but not synonymous substitutions and so attributed their co-occurrence to functional constraints (Callahan *et al.*, 2011). While this does not seem to be the case for the synonymous mutations in our analysis, together these results suggest that physical linkage may often be important in generating signals of correlated evolution. If so, this result suggests using caution when interpreting the results of computational studies that identify strong correlated evolution among traits or sites within a genome.

While computational methods provide us with a high-throughput means of determining candidates for epistasis, we often lack a sufficient understanding of epistasis at the molecular level to have confidence in this interpretation in the absence of additional evidence. Our work represents a first step in this direction: by using a computational approach to predict candidates for correlated evolution, with follow-up analyses suggesting that only a subset of these candidates are actually under epistasis, we were able to provide direct experimental evidence that epistasis underlies at least some instances of correlated evolution. However, this interpretation relies on our ability to conduct experimental validations under selective conditions that we presume to closely resemble those in which the sites evolved. This may not always be possible. Indeed, most of the strongest instances of correlated evolution in our data set appear to be unconnected to fitness or epistasis in an antibiotic selective context. As densely sampled population genomic data are not always available (such as the *>* 3000 genomes in Skwark *et al.* (2017)), we urge caution in interpreting instances of correlation between sites from high throughput computational analyses as solely due to epistasis.

## ACKNOWLEDGEMENTS

We would like to thank Felipe Dargent and Matti Ruuskanen, as well as three anonymous reviewers, for their constructive feedbacks on the manuscript. We are grateful for the computational resources of the Centre for Advanced Computing at Queens University. This work was funded by the Natural Sciences and Engineering Research Council of Canada (S.A.B. and R.K.).

## DATA ACCESSIBILITY

All R scripts and complete results of the computational analysis of correlated evolution can be found at github.com/JDench/sigResults_AEGIS_inVivo.

