## Supplementary Information 1 for "Identifying the drivers of computationally detected correlated evolution among sites under antibiotic selection"

### 1 Supplementary Text

#### 1.1 Additional Methods Details

**Alignment algorithm** We developed a custom algorithm to reconstruct an genome alignment, made of concatenated genes (ie: an exome alignment), using as reference a PA14 genome database obtained from `www.pseudomonas.com` (Winsor *et al.*, 2016, accessed Dec 4, 2014). From this database, we extracted the nucleotide sequences for the 5,977 genes, and used each of them as a query for BLASTn (Altschul *et al.*, 1990) searches against a local database containing the 390 draft genomes, as well as the three complete genomes of PA01, PA7 and PA14. Results with more than 90% identity were then used to construct scaffolds, which were subsequently extended, for each of the 390 draft genomes, making sure that the same genomic information was not used more than once. The extended scaffold sequences of each gene were aligned with MUSCLE (Edgar, 2004). We discarded genes present in less than 50% of strains with $\geq$ 90% of the length of the PA14 query (non-gap characters). The remaining genes were concatenated into a single exome alignment with *catfasta2phyml.pl* (Nylander, 2015).

**Complexity Hypothesis** The species tree for our exome alignment was reconstructed based on highly networked genes that, according to the complexity hypothesis, are unlikely to be horizontally transferred. According to the complexity hypothesis (Jain *et al.*, 1999; Aris-Brosou, 2005), highly networked genes are those whose products are involved in complex multiprotein interactions such as information processing genes (e.g. rRNA genes, tRNA genes, transcription and translation polymerases). Based on the functional annotations of PA14 genes obtained from the Clusters of Orthologous Groups database (Tatusov *et al.*, 1997; Galperin *et al.*, 2015), we extracted 1,290 information genes (those with COG terms A, B, J, K, and L), and concatenated them into a single alignment used to estimate the species tree.

**Analysis of correlated evolution** Recoding our concatenated exome into binary states resulted in 410,665 polymorphic sites. We estimated that a full pairwise comparison of the polymorphic sites would have required more than one year of computing even when parallelized across twelve computational nodes each with four processors. It is for this reason that we performed a 12 gene by exome analysis.

**Modified selection counter-selection protocol:** Our modified WT allele replacement protocol is based on a selection counter-selection method (Schweizer, 2008), whereby a vector-borne mutant allele recombines with homologous chromosomal sequences before the delivery vector is itself removed from the targeted genome. Mutant alleles of *gyrA* and *parC* were generated from WT chromosomal DNA by amplification of each locus in paired PCR reactions that overlapped at sequences adjacent to the introduced substitution. One of the overlapping primers coded for the substitution of interest through a mismatch at the target site. The paired amplicons were ligated to an allelic replacement vector using Golden Gate assembly (Engler *et al.*, 2008), which permitted scarless ligation of the PCR products (see Table S2 for details). The vector, derived from pAH79 (Melnyk *et al.*, 2017) and modified for Golden Gate cloning, includes the $TetA$ selectable marker and $sacB$ counter-selection gene. After ligation, vectors were transformed into chemically competent *Escherichia coli* (DH5 $\lambda$pir). The mutant alleles were transferred into *P. aeruginosa* strains as previously described (Melnyk *et al.*, 2017) via tri-parental conjugation, involving a helper *E. coli* that carries pRK2013 (Figurski and Helinski, 1979). Through a round of selection (LB agar with 100$\mu$g/mL Nitrofurantoin and 80$\mu$g/mL Tetracycline), we isolated colonies whose genome had recombined with the mutant allele. The recombinants were subsequently counter-selected (LB agar with 5% sucrose) for loss of the plasmid sequences. Mutant constructs were first confirmed by sequencing the region targeted for replacement. To identify constructs least likely to contain secondary mutations outside the sequenced region, we performed competitive fitness assays with a minimum of four independent constructs, and accepted constructs with relatively similar fitness measures for downstream analyses.

**Calculating competitive fitness:** Competitive fitness was calculated as $\omega = (f_{final} - f_{initial})^{(1/generations)}$, where $f_{initial}$ and $f_{final}$ are the initial and final frequency of focal strains, and the number of generations was based on the dilution factor, calculated as $\log_2(100) \sim 6.64$. The evidence for epistasis ($\epsilon$) was calculated with a multiplicative fitness model (Trindade *et al.*, 2009) such that $\epsilon = W_{WT}W_{AB} - W_A W_B$ where W stands for fitness and the subscripts (WT, A, B, AB) represent the wild-type, single and double mutant genotypes. There is evidence for epistasis when the $\epsilon$ value is greater than our measurement error estimated via error propagation.

**Adaptive boosting algorithm** Adaptive boosting is a supervised machine learning algorithm that uses the weighted sum of many sequentially fitted classifiers, and is considered the "best out-of-the-box" classifier in part because it yields lower classification error rates while being less susceptible to overfitting (James *et al.*, 2013). Nucleotide sites in our exome alignment were quantified for their importance in predicting levofloxacin resistance by computing the number of fitted classification trees which retain a predictor, in this case nucleotide site. Classification was based on the association with the discrete phenotype of sensitivity or resistance to levofloxacin. Phenotype data were obtained from the original study that published the 390 draft genomes used to construct our alignment (Kos *et al.*, 2015).

**Calculating $\Delta G$** To estimate biological effects of synonymous mutations, $\Delta G$ values were calculated with the mFold server (Zuker, 2003), based on the QuickFold service with default settings (`unafold.rna.albany.edu/?q=DINAMelt/Quickfold`). Similar to previous studies (Takanami and Zubay, 1964), we calculated the free energy of each mRNA transcript by considering 50 nucleotides upstream and downstream of each substitution. Mean and error values of $\Delta G_{rel}$ were calculated using weighted values for all the predicted folding structures returned by QuickFold.

**Calculating $I_{TE}$** Also, to estimate biological effects of synonymous mutations, we calculated the index of translation elongation ($I_{TE}$: Xia, 2014) with default settings in DAMBE ver. 6

(Xia, 2017). These calculations used the codon frequency of genes that are highly expressed in *P. aeruginosa* (Hilterbrand *et al.*, 2012), and lowly expressed genes calculated using all of the other genes in our alignment. Files containing the codon frequency counts for all strains in our alignment are available from `github.com/JDench/Pseudomonas_DAMBE_ITE`.

#### 1.2 Supplementary analyses

While a previous study has shown the excellent specificity of AEGIS at identifying correlated pairs of substitutions from simulated evolution (Nshogozabahizi *et al.*, 2017), the $\approx 127,000$ significantly correlated pairs ($P \leq 0.01$) was much higher than expected and we deemed it beneficial to review the results prior to *in vitro* study. As this study is the first to have tried to identify correlated evolution among pairs of sites in the whole genome of *P. aeruginosa* there was no general dataset against which we could compare our results, instead we performed summary analyses to describe trends in the results. Recall that for reasons of computational time, this study analysed the evidence of correlated evolution between the sites in 12 genes and the rest of the genome. If the results of AEGIS were largely false positives, we would have expected the number of polymorphic sites (*i.e.* sites in our alignment with different nucleotide characters across genotypes) in a gene to be positively correlated with the number of correlated pairs involving substitutions in that gene. We found no evidence that the number of polymorphic sites in the 12 genes was correlated to the number of associated significant pairs (Pearson's $\rho$, $t = 0.5530$, $df = 10$, $P = 0.5924$). Further, we found no evidence that the proportion of polymorphic sites in a gene (i.e., number of sites divided by gene length) differed between the six focal genes ($gyrA$, $gyrB$, $morA$, $nfxB$, $parC$, $parE$), the additional six genes chosen at random ($dnaA$, $dnaN$, $lpd3$, $ribD$, $rpoB$, $serC$), and any of the other genes in our alignment (ANOVA, $F = 0.1227$, $df = 2$, $P = 0.8845$). This led us to conclude that the results of AEGIS reflected some true evolutionary signal.

While we did not have specific information concerning the *in vitro* effects of most substitutions identified in the results, we could define several descriptive traits. For this, we compared trends among our significantly correlated paired sites to what we would expect by chance (*i.e.* from a null model). If chance alone drove our detection of correlated site pairs, we would expect to detect a paired traits according to their frequency in our exome alignment. We found that the frequency of observed paired traits differs from our null expectation (Fig. S1). As a first assessment we leveraged our *a priori* assumption that mutations in the six focal genes were more likely to show an adaptive response to fluoroquinolone selection compared to the other six genes. If our assumption was correct and the results of AEGIS reflect correlated evolution in response to selection, we would expect more pairs to include mutations in the six focal genes. When comparing the number of correlated pairs which include zero, one, or two sites in the six focal genes, we found that pairs with at least one site in an expected gene are consistently higher than by chance (compare panels a,b in Fig. S1). We next wanted to assess if correlated pairs of potentially adaptive substitutions (i.e. nonsynonymous) were detected more often than by chance. While synonymous substitutions may be selected for (Agashe *et al.*, 2016; Bailey *et al.*, 2014), in response to antibiotic selection we assumed that only nonsynonymous substitutions were likely to be adaptive. We looked for differences in the expected and observed number of correlated pairs where zero, one or two sites were nonsynonymous (compare Fig. S1 c,d). We found that nonsynonymous pairs are dramatically underrepresented comprising only 2 pairs with at least medium (*parC* 786 and PA14_34000 967 - hypothetical type VI secretion protein - $10^{-7} \leq P \leq 10^{-6}$) and strong ($gyrA$ 248 and *parC* 260, $P \leq 10^{-11}$) evidence for correlated evolution respectively. We interpret the higher than expected number of synonymous pairs to suggest that hitchhiking (Maynard Smith and Haigh, 1974), possibly as “cohorts” (Lang *et al.*, 2013), explains the majority of correlated pairs identified by AEGIS.

### SI Tables

**Table S1. Table of strains included in whole exome alignment.** Where known we provide the name of isolates used in our study along with the year, country and origin as per the cited reference paper.

| Isolate | Year of isolation | Country | City | Reference |
|---|---|---|---|---|
| AZPAE12135 | 2005 | United States | New York | (Kos *et al.*, 2015) |
| AZPAE12136 | 2005 | United States | New York | (Kos *et al.*, 2015) |
| AZPAE12137 | 2005 | United States | New York | (Kos *et al.*, 2015) |
| AZPAE12138 | 2005 | United States | New York | (Kos *et al.*, 2015) |
| AZPAE12140 | 2005 | United States | New York | (Kos *et al.*, 2015) |
| AZPAE12142 | 2005 | United States | New York | (Kos *et al.*, 2015) |
| AZPAE12143 | 2005 | United States | New York | (Kos *et al.*, 2015) |
| AZPAE12144 | 2005 | United States | New York | (Kos *et al.*, 2015) |
| AZPAE12145 | 2005 | United States | New York | (Kos *et al.*, 2015) |
| AZPAE12146 | 2005 | United States | New York | (Kos *et al.*, 2015) |
| AZPAE12147 | 2005 | United States | New York | (Kos *et al.*, 2015) |
| AZPAE12148 | 2005 | United States | New York | (Kos *et al.*, 2015) |
| AZPAE12149 | 2005 | United States | New York | (Kos *et al.*, 2015) |
| AZPAE12150 | 2005 | United States | New York | (Kos *et al.*, 2015) |
| AZPAE12151 | 2005 | United States | New York | (Kos *et al.*, 2015) |
| AZPAE12152 | 2005 | United States | New York | (Kos *et al.*, 2015) |
| AZPAE12153 | 2005 | United States | New York | (Kos *et al.*, 2015) |
| AZPAE12154 | 2005 | United States | New York | (Kos *et al.*, 2015) |
| AZPAE12155 | 2005 | United States | New York | (Kos *et al.*, 2015) |
| AZPAE12156 | 2005 | United States | New York | (Kos *et al.*, 2015) |
| AZPAE12409 | 2007 | United States | Cleveland | (Kos *et al.*, 2015) |
| AZPAE12410 | 2007 | United States | Cleveland | (Kos *et al.*, 2015) |
| AZPAE12411 | 2007 | United States | Cleveland | (Kos *et al.*, 2015) |
| AZPAE12412 | 2007 | United States | Cleveland | (Kos *et al.*, 2015) |
| AZPAE12413 | 2007 | United States | Cleveland | (Kos *et al.*, 2015) |
| AZPAE12414 | 2007 | United States | Cleveland | (Kos *et al.*, 2015) |
| AZPAE12415 | 2007 | United States | Cleveland | (Kos *et al.*, 2015) |
| AZPAE12416 | 2007 | United States | Cleveland | (Kos *et al.*, 2015) |
| AZPAE12417 | 2007 | United States | Cleveland | (Kos *et al.*, 2015) |
| AZPAE12418 | 2007 | United States | Cleveland | (Kos *et al.*, 2015) |
| AZPAE12419 | 2007 | United States | Cleveland | (Kos *et al.*, 2015) |
| AZPAE12420 | 2007 | United States | Cleveland | (Kos *et al.*, 2015) |
| AZPAE12421 | 2007 | United States | Cleveland | (Kos *et al.*, 2015) |

Continued on next page

**Table S1 – continued from previous page**

| Isolat | Year of isolation | Country | City | Reference |
|---|---|---|---|---|
| AZPAE12422 | 2007 | United States | Cleveland | (Kos *et al.*, 2015) |
| AZPAE12423 | 2007 | United States | Cleveland | (Kos *et al.*, 2015) |
| AZPAE13756 | 2009 | Canada | unknown | (Kos *et al.*, 2015) |
| AZPAE13757 | 2009 | Canada | unknown | (Kos *et al.*, 2015) |
| AZPAE13848 | 2010 | India | unknown | (Kos *et al.*, 2015) |
| AZPAE13850 | 2010 | India | unknown | (Kos *et al.*, 2015) |
| AZPAE13853 | 2010 | India | unknown | (Kos *et al.*, 2015) |
| AZPAE13856 | 2010 | India | unknown | (Kos *et al.*, 2015) |
| AZPAE13858 | 2010 | India | unknown | (Kos *et al.*, 2015) |
| AZPAE13860 | 2010 | India | unknown | (Kos *et al.*, 2015) |
| AZPAE13864 | 2010 | India | unknown | (Kos *et al.*, 2015) |
| AZPAE13866 | 2010 | China | unknown | (Kos *et al.*, 2015) |
| AZPAE13872 | 2010 | Mexico | unknown | (Kos *et al.*, 2015) |
| AZPAE13876 | 2010 | Portugal | unknown | (Kos *et al.*, 2015) |
| AZPAE13877 | 2010 | Romania | unknown | (Kos *et al.*, 2015) |
| AZPAE13879 | 2010 | Argentina | unknown | (Kos *et al.*, 2015) |
| AZPAE13880 | 2010 | Mexico | unknown | (Kos *et al.*, 2015) |
| AZPAE14352 | 2010 | France | unknown | (Kos *et al.*, 2015) |
| AZPAE14353 | 2010 | France | unknown | (Kos *et al.*, 2015) |
| AZPAE14359 | 2010 | China | Shatin | (Kos *et al.*, 2015) |
| AZPAE14372 | 2010 | China | Hong Kong | (Kos *et al.*, 2015) |
| AZPAE14373 | 2010 | Germany | München | (Kos *et al.*, 2015) |
| AZPAE14379 | 2010 | Germany | Heidelberg | (Kos *et al.*, 2015) |
| AZPAE14381 | 2010 | Spain | Bilbao | (Kos *et al.*, 2015) |
| AZPAE14390 | 2011 | China | Hong Kong | (Kos *et al.*, 2015) |
| AZPAE14393 | 2011 | Spain | Madrid | (Kos *et al.*, 2015) |
| AZPAE14394 | 2011 | Spain | Madrid | (Kos *et al.*, 2015) |
| AZPAE14395 | 2011 | Spain | Bilbao | (Kos *et al.*, 2015) |
| AZPAE14398 | 2011 | Germany | München | (Kos *et al.*, 2015) |
| AZPAE14402 | 2011 | France | Rouen | (Kos *et al.*, 2015) |
| AZPAE14403 | 2011 | France | Rouen | (Kos *et al.*, 2015) |
| AZPAE14404 | 2012 | China | Shatin | (Kos *et al.*, 2015) |
| AZPAE14410 | 2012 | Germany | Heidelberg | (Kos *et al.*, 2015) |
| AZPAE14415 | 2009 | Portugal | unknown | (Kos *et al.*, 2015) |
| AZPAE14422 | 2009 | United States | Roseburg | (Kos *et al.*, 2015) |
| AZPAE14437 | 2010 | Canada | unknown | (Kos *et al.*, 2015) |
| AZPAE14441 | 2010 | Taiwan | unknown | (Kos *et al.*, 2015) |
| AZPAE14442 | 2010 | Taiwan | unknown | (Kos *et al.*, 2015) |

Continued on next page

**Table S1 – continued from previous page**

| Isolat | Year of isolation | Country | City | Reference |
|---|---|---|---|---|
| AZPAE14443 | 2010 | United States | unknown | (Kos *et al.*, 2015) |
| AZPAE14453 | 2011 | United States | Detroit | (Kos *et al.*, 2015) |
| AZPAE14463 | 2011 | Colombia | Bogota | (Kos *et al.*, 2015) |
| AZPAE14499 | 2011 | Spain | Santander | (Kos *et al.*, 2015) |
| AZPAE14505 | 2011 | France | Paris | (Kos *et al.*, 2015) |
| AZPAE14509 | 2011 | France | Nantes | (Kos *et al.*, 2015) |
| AZPAE14526 | 2010 | Spain | Palma de Mallorca | (Kos *et al.*, 2015) |
| AZPAE14533 | 2011 | Germany | Koln | (Kos *et al.*, 2015) |
| AZPAE14535 | 2010 | Spain | Santander | (Kos *et al.*, 2015) |
| AZPAE14538 | 2010 | China | Shatin | (Kos *et al.*, 2015) |
| AZPAE14550 | 2010 | China | Beijing | (Kos *et al.*, 2015) |
| AZPAE14554 | 2010 | Spain | Santander | (Kos *et al.*, 2015) |
| AZPAE14557 | 2010 | Germany | Koln | (Kos *et al.*, 2015) |
| AZPAE14566 | 2011 | China | Shatin | (Kos *et al.*, 2015) |
| AZPAE14570 | 2010 | Germany | unknown | (Kos *et al.*, 2015) |
| AZPAE14687 | 2012 | Mexico | unknown | (Kos *et al.*, 2015) |
| AZPAE14688 | 2012 | Mexico | unknown | (Kos *et al.*, 2015) |
| AZPAE14689 | 2012 | Mexico | unknown | (Kos *et al.*, 2015) |
| AZPAE14690 | 2012 | Romania | unknown | (Kos *et al.*, 2015) |
| AZPAE14691 | 2012 | United States | unknown | (Kos *et al.*, 2015) |
| AZPAE14692 | 2012 | United States | unknown | (Kos *et al.*, 2015) |
| AZPAE14693 | 2012 | Romania | unknown | (Kos *et al.*, 2015) |
| AZPAE14694 | 2012 | Romania | unknown | (Kos *et al.*, 2015) |
| AZPAE14695 | 2012 | Israel | unknown | (Kos *et al.*, 2015) |
| AZPAE14697 | 2012 | Israel | unknown | (Kos *et al.*, 2015) |
| AZPAE14698 | 2012 | Israel | unknown | (Kos *et al.*, 2015) |
| AZPAE14699 | 2012 | United States | unknown | (Kos *et al.*, 2015) |
| AZPAE14700 | 2012 | Philippines | unknown | (Kos *et al.*, 2015) |
| AZPAE14701 | 2012 | Philippines | unknown | (Kos *et al.*, 2015) |
| AZPAE14702 | 2012 | Philippines | unknown | (Kos *et al.*, 2015) |
| AZPAE14703 | 2012 | Philippines | unknown | (Kos *et al.*, 2015) |
| AZPAE14704 | 2012 | Greece | unknown | (Kos *et al.*, 2015) |
| AZPAE14705 | 2012 | Greece | unknown | (Kos *et al.*, 2015) |
| AZPAE14706 | 2012 | Greece | unknown | (Kos *et al.*, 2015) |
| AZPAE14707 | 2012 | Greece | unknown | (Kos *et al.*, 2015) |
| AZPAE14708 | 2012 | Greece | unknown | (Kos *et al.*, 2015) |
| AZPAE14710 | 2012 | United States | unknown | (Kos *et al.*, 2015) |
| AZPAE14711 | 2012 | Venezuela | unknown | (Kos *et al.*, 2015) |

Continued on next page

**Table S1 – continued from previous page**

| Isolat | Year of isolation | Country | City | Reference |
|---|---|---|---|---|
| AZPAE14712 | 2012 | Venezuela | unknown | (Kos *et al.*, 2015) |
| AZPAE14713 | 2012 | Venezuela | unknown | (Kos *et al.*, 2015) |
| AZPAE14714 | 2012 | Venezuela | unknown | (Kos *et al.*, 2015) |
| AZPAE14715 | 2012 | Venezuela | unknown | (Kos *et al.*, 2015) |
| AZPAE14716 | 2012 | Venezuela | unknown | (Kos *et al.*, 2015) |
| AZPAE14717 | 2012 | United States | unknown | (Kos *et al.*, 2015) |
| AZPAE14718 | 2012 | United States | unknown | (Kos *et al.*, 2015) |
| AZPAE14719 | 2012 | Colombia | unknown | (Kos *et al.*, 2015) |
| AZPAE14720 | 2012 | Colombia | unknown | (Kos *et al.*, 2015) |
| AZPAE14721 | 2012 | Colombia | unknown | (Kos *et al.*, 2015) |
| AZPAE14722 | 2012 | Italy | unknown | (Kos *et al.*, 2015) |
| AZPAE14723 | 2012 | Italy | unknown | (Kos *et al.*, 2015) |
| AZPAE14724 | 2012 | Italy | unknown | (Kos *et al.*, 2015) |
| AZPAE14725 | 2012 | United States | unknown | (Kos *et al.*, 2015) |
| AZPAE14726 | 2012 | United States | unknown | (Kos *et al.*, 2015) |
| AZPAE14727 | 2012 | United States | unknown | (Kos *et al.*, 2015) |
| AZPAE14728 | 2012 | United States | unknown | (Kos *et al.*, 2015) |
| AZPAE14729 | 2012 | Italy | unknown | (Kos *et al.*, 2015) |
| AZPAE14730 | 2012 | Italy | unknown | (Kos *et al.*, 2015) |
| AZPAE14731 | 2012 | Italy | unknown | (Kos *et al.*, 2015) |
| AZPAE14732 | 2012 | United States | unknown | (Kos *et al.*, 2015) |
| AZPAE14809 | 2004 | India | Mumbai | (Kos *et al.*, 2015) |
| AZPAE14810 | 2004 | India | Mumbai | (Kos *et al.*, 2015) |
| AZPAE14811 | 2004 | India | Mumbai | (Kos *et al.*, 2015) |
| AZPAE14812 | 2004 | India | Mumbai | (Kos *et al.*, 2015) |
| AZPAE14813 | 2004 | India | Mumbai | (Kos *et al.*, 2015) |
| AZPAE14814 | 2004 | France | Besancon | (Kos *et al.*, 2015) |
| AZPAE14815 | 2004 | France | Besancon | (Kos *et al.*, 2015) |
| AZPAE14816 | 2004 | France | Besancon | (Kos *et al.*, 2015) |
| AZPAE14817 | 2004 | France | Besancon | (Kos *et al.*, 2015) |
| AZPAE14818 | 2004 | France | Besancon | (Kos *et al.*, 2015) |
| AZPAE14819 | 2004 | Brazil | Sao Paulo | (Kos *et al.*, 2015) |
| AZPAE14820 | 2004 | Brazil | Sao Paulo | (Kos *et al.*, 2015) |
| AZPAE14821 | 2004 | Brazil | Sao Paulo | (Kos *et al.*, 2015) |
| AZPAE14822 | 2004 | Brazil | Sao Paulo | (Kos *et al.*, 2015) |
| AZPAE14823 | 2005 | Germany | Koln | (Kos *et al.*, 2015) |
| AZPAE14824 | 2005 | Germany | Koln | (Kos *et al.*, 2015) |
| AZPAE14825 | 2005 | Germany | Koln | (Kos *et al.*, 2015) |

Continued on next page

**Table S1 – continued from previous page**

| Isolat | Year of isolation | Country | City | Reference |
|---|---|---|---|---|
| AZPAE14826 | 2005 | United States | Detroit | (Kos *et al.*, 2015) |
| AZPAE14827 | 2005 | United States | Detroit | (Kos *et al.*, 2015) |
| AZPAE14828 | 2005 | United States | Detroit | (Kos *et al.*, 2015) |
| AZPAE14829 | 2005 | United States | Detroit | (Kos *et al.*, 2015) |
| AZPAE14830 | 2005 | Argentina | Victoria | (Kos *et al.*, 2015) |
| AZPAE14831 | 2005 | Argentina | Victoria | (Kos *et al.*, 2015) |
| AZPAE14832 | 2005 | Argentina | Victoria | (Kos *et al.*, 2015) |
| AZPAE14833 | 2005 | Argentina | Victoria | (Kos *et al.*, 2015) |
| AZPAE14834 | 2005 | Argentina | Victoria | (Kos *et al.*, 2015) |
| AZPAE14835 | 2006 | China | Beijing | (Kos *et al.*, 2015) |
| AZPAE14836 | 2006 | China | Beijing | (Kos *et al.*, 2015) |
| AZPAE14837 | 2006 | China | Beijing | (Kos *et al.*, 2015) |
| AZPAE14838 | 2006 | China | Beijing | (Kos *et al.*, 2015) |
| AZPAE14839 | 2006 | China | Beijing | (Kos *et al.*, 2015) |
| AZPAE14840 | 2006 | China | Beijing | (Kos *et al.*, 2015) |
| AZPAE14841 | 2006 | China | Beijing | (Kos *et al.*, 2015) |
| AZPAE14842 | 2006 | United States | Roseburg | (Kos *et al.*, 2015) |
| AZPAE14843 | 2006 | United States | Roseburg | (Kos *et al.*, 2015) |
| AZPAE14844 | 2006 | United States | Roseburg | (Kos *et al.*, 2015) |
| AZPAE14845 | 2006 | Germany | Koln | (Kos *et al.*, 2015) |
| AZPAE14846 | 2006 | France | Nantes | (Kos *et al.*, 2015) |
| AZPAE14847 | 2006 | France | Nantes | (Kos *et al.*, 2015) |
| AZPAE14848 | 2006 | France | Nantes | (Kos *et al.*, 2015) |
| AZPAE14850 | 2006 | France | Nantes | (Kos *et al.*, 2015) |
| AZPAE14851 | 2006 | France | Nantes | (Kos *et al.*, 2015) |
| AZPAE14852 | 2005 | Brazil | Sao Paulo | (Kos *et al.*, 2015) |
| AZPAE14853 | 2007 | Brazil | Curitiba | (Kos *et al.*, 2015) |
| AZPAE14855 | 2007 | France | Paris | (Kos *et al.*, 2015) |
| AZPAE14856 | 2007 | France | Paris | (Kos *et al.*, 2015) |
| AZPAE14857 | 2007 | France | Paris | (Kos *et al.*, 2015) |
| AZPAE14858 | 2007 | France | Paris | (Kos *et al.*, 2015) |
| AZPAE14859 | 2007 | Spain | Bilbao | (Kos *et al.*, 2015) |
| AZPAE14860 | 2007 | Spain | Bilbao | (Kos *et al.*, 2015) |
| AZPAE14861 | 2007 | Spain | Bilbao | (Kos *et al.*, 2015) |
| AZPAE14862 | 2007 | India | Chennai | (Kos *et al.*, 2015) |
| AZPAE14863 | 2007 | India | Chennai | (Kos *et al.*, 2015) |
| AZPAE14864 | 2007 | India | Chennai | (Kos *et al.*, 2015) |
| AZPAE14865 | 2007 | India | Chennai | (Kos *et al.*, 2015) |

Continued on next page

**Table S1 – continued from previous page**

| Isolat | Year of isolation | Country | City | Reference |
|---|---|---|---|---|
| AZPAE14866 | 2007 | China | Shatin | (Kos *et al.*, 2015) |
| AZPAE14867 | 2007 | China | Shatin | (Kos *et al.*, 2015) |
| AZPAE14868 | 2007 | Argentina | Victoria | (Kos *et al.*, 2015) |
| AZPAE14869 | 2007 | Argentina | Victoria | (Kos *et al.*, 2015) |
| AZPAE14870 | 2007 | Argentina | Victoria | (Kos *et al.*, 2015) |
| AZPAE14871 | 2007 | Argentina | Victoria | (Kos *et al.*, 2015) |
| AZPAE14872 | 2007 | Argentina | Victoria | (Kos *et al.*, 2015) |
| AZPAE14873 | 2007 | Argentina | Victoria | (Kos *et al.*, 2015) |
| AZPAE14874 | 2007 | United States | Detroit | (Kos *et al.*, 2015) |
| AZPAE14875 | 2007 | United States | Detroit | (Kos *et al.*, 2015) |
| AZPAE14876 | 2007 | United States | Detroit | (Kos *et al.*, 2015) |
| AZPAE14877 | 2007 | United States | Roseburg | (Kos *et al.*, 2015) |
| AZPAE14878 | 2007 | United States | Roseburg | (Kos *et al.*, 2015) |
| AZPAE14879 | 2007 | United States | Roseburg | (Kos *et al.*, 2015) |
| AZPAE14880 | 2007 | Spain | Santander | (Kos *et al.*, 2015) |
| AZPAE14881 | 2007 | Spain | Santander | (Kos *et al.*, 2015) |
| AZPAE14882 | 2007 | Spain | Santander | (Kos *et al.*, 2015) |
| AZPAE14883 | 2007 | Croatia | Split | (Kos *et al.*, 2015) |
| AZPAE14884 | 2007 | Croatia | Split | (Kos *et al.*, 2015) |
| AZPAE14885 | 2007 | Croatia | Split | (Kos *et al.*, 2015) |
| AZPAE14886 | 2007 | Croatia | Split | (Kos *et al.*, 2015) |
| AZPAE14887 | 2007 | Croatia | Split | (Kos *et al.*, 2015) |
| AZPAE14888 | 2007 | United States | Detroit | (Kos *et al.*, 2015) |
| AZPAE14889 | 2008 | China | Shatin | (Kos *et al.*, 2015) |
| AZPAE14890 | 2008 | France | Besancon | (Kos *et al.*, 2015) |
| AZPAE14891 | 2008 | France | Besancon | (Kos *et al.*, 2015) |
| AZPAE14892 | 2008 | France | Besancon | (Kos *et al.*, 2015) |
| AZPAE14893 | 2008 | France | Besancon | (Kos *et al.*, 2015) |
| AZPAE14894 | 2008 | Germany | Heidelberg | (Kos *et al.*, 2015) |
| AZPAE14895 | 2008 | Germany | Heidelberg | (Kos *et al.*, 2015) |
| AZPAE14897 | 2008 | India | Chennai | (Kos *et al.*, 2015) |
| AZPAE14898 | 2008 | India | Chennai | (Kos *et al.*, 2015) |
| AZPAE14899 | 2008 | India | Chennai | (Kos *et al.*, 2015) |
| AZPAE14900 | 2008 | India | Chennai | (Kos *et al.*, 2015) |
| AZPAE14901 | 2008 | India | Chennai | (Kos *et al.*, 2015) |
| AZPAE14902 | 2008 | Argentina | Victoria | (Kos *et al.*, 2015) |
| AZPAE14903 | 2008 | Spain | Madrid | (Kos *et al.*, 2015) |
| AZPAE14904 | 2008 | Spain | Madrid | (Kos *et al.*, 2015) |

Continued on next page

**Table S1 – continued from previous page**

| Isolat | Year of isolation | Country | City | Reference |
|---|---|---|---|---|
| AZPAE14905 | 2008 | Germany | Koln | (Kos *et al.*, 2015) |
| AZPAE14906 | 2008 | Germany | Koln | (Kos *et al.*, 2015) |
| AZPAE14907 | 2008 | China | Shatin | (Kos *et al.*, 2015) |
| AZPAE14908 | 2008 | Spain | Palma de Mallorca | (Kos *et al.*, 2015) |
| AZPAE14909 | 2008 | Spain | Palma de Mallorca | (Kos *et al.*, 2015) |
| AZPAE14910 | 2008 | India | Mumbai | (Kos *et al.*, 2015) |
| AZPAE14911 | 2008 | India | Mumbai | (Kos *et al.*, 2015) |
| AZPAE14912 | 2008 | Croatia | Split | (Kos *et al.*, 2015) |
| AZPAE14913 | 2008 | Croatia | Split | (Kos *et al.*, 2015) |
| AZPAE14914 | 2008 | Spain | Santander | (Kos *et al.*, 2015) |
| AZPAE14915 | 2008 | Spain | Santander | (Kos *et al.*, 2015) |
| AZPAE14916 | 2008 | Spain | Bilbao | (Kos *et al.*, 2015) |
| AZPAE14917 | 2008 | Spain | Bilbao | (Kos *et al.*, 2015) |
| AZPAE14918 | 2008 | Spain | Bilbao | (Kos *et al.*, 2015) |
| AZPAE14919 | 2008 | Spain | Bilbao | (Kos *et al.*, 2015) |
| AZPAE14920 | 2008 | Spain | Bilbao | (Kos *et al.*, 2015) |
| AZPAE14921 | 2009 | France | Paris | (Kos *et al.*, 2015) |
| AZPAE14922 | 2009 | France | Paris | (Kos *et al.*, 2015) |
| AZPAE14923 | 2008 | Brazil | Sao Paulo | (Kos *et al.*, 2015) |
| AZPAE14924 | 2008 | Brazil | Sao Paulo | (Kos *et al.*, 2015) |
| AZPAE14925 | 2008 | Brazil | Sao Paulo | (Kos *et al.*, 2015) |
| AZPAE14926 | 2008 | Brazil | Sao Paulo | (Kos *et al.*, 2015) |
| AZPAE14927 | 2008 | Brazil | Sao Paulo | (Kos *et al.*, 2015) |
| AZPAE14928 | 2008 | Brazil | Sao Paulo | (Kos *et al.*, 2015) |
| AZPAE14929 | 2009 | Germany | Aachen | (Kos *et al.*, 2015) |
| AZPAE14930 | 2009 | Germany | Aachen | (Kos *et al.*, 2015) |
| AZPAE14931 | 2009 | Germany | Aachen | (Kos *et al.*, 2015) |
| AZPAE14932 | 2009 | Germany | Aachen | (Kos *et al.*, 2015) |
| AZPAE14933 | 2009 | France | Rouen | (Kos *et al.*, 2015) |
| AZPAE14934 | 2009 | France | Rouen | (Kos *et al.*, 2015) |
| AZPAE14935 | 2008 | France | Rouen | (Kos *et al.*, 2015) |
| AZPAE14936 | 2008 | Brazil | Curitiba | (Kos *et al.*, 2015) |
| AZPAE14937 | 2008 | France | Rouen | (Kos *et al.*, 2015) |
| AZPAE14938 | 2008 | France | Rouen | (Kos *et al.*, 2015) |
| AZPAE14939 | 2009 | Colombia | Bogota | (Kos *et al.*, 2015) |
| AZPAE14940 | 2009 | France | Rouen | (Kos *et al.*, 2015) |
| AZPAE14941 | 2009 | China | Hong Kong | (Kos *et al.*, 2015) |
| AZPAE14942 | 2009 | China | Hong Kong | (Kos *et al.*, 2015) |

Continued on next page

**Table S1 – continued from previous page**

| Isolat | Year of isolation | Country | City | Reference |
|---|---|---|---|---|
| AZPAE14943 | 2009 | Colombia | Bogota | (Kos *et al.*, 2015) |
| AZPAE14944 | 2009 | United States | Detroit | (Kos *et al.*, 2015) |
| AZPAE14945 | 2009 | United States | Detroit | (Kos *et al.*, 2015) |
| AZPAE14946 | 2009 | United States | Detroit | (Kos *et al.*, 2015) |
| AZPAE14947 | 2009 | China | Beijing | (Kos *et al.*, 2015) |
| AZPAE14948 | 2009 | Argentina | Victoria | (Kos *et al.*, 2015) |
| AZPAE14949 | 2009 | Argentina | Victoria | (Kos *et al.*, 2015) |
| AZPAE14950 | 2009 | Argentina | Victoria | (Kos *et al.*, 2015) |
| AZPAE14951 | 2009 | Argentina | Victoria | (Kos *et al.*, 2015) |
| AZPAE14952 | 2009 | China | Shatin | (Kos *et al.*, 2015) |
| AZPAE14953 | 2009 | China | Shatin | (Kos *et al.*, 2015) |
| AZPAE14954 | 2009 | France | Nantes | (Kos *et al.*, 2015) |
| AZPAE14955 | 2009 | France | Nantes | (Kos *et al.*, 2015) |
| AZPAE14956 | 2009 | Germany | München | (Kos *et al.*, 2015) |
| AZPAE14957 | 2009 | Germany | München | (Kos *et al.*, 2015) |
| AZPAE14958 | 2009 | India | Mumbai | (Kos *et al.*, 2015) |
| AZPAE14959 | 2009 | India | Mumbai | (Kos *et al.*, 2015) |
| AZPAE14960 | 2009 | Spain | Palma de Mallorca | (Kos *et al.*, 2015) |
| AZPAE14961 | 2009 | Spain | Palma de Mallorca | (Kos *et al.*, 2015) |
| AZPAE14962 | 2009 | Spain | Palma de Mallorca | (Kos *et al.*, 2015) |
| AZPAE14963 | 2009 | Spain | Palma de Mallorca | (Kos *et al.*, 2015) |
| AZPAE14964 | 2009 | France | Nantes | (Kos *et al.*, 2015) |
| AZPAE14965 | 2009 | France | Nantes | (Kos *et al.*, 2015) |
| AZPAE14967 | 2009 | Croatia | Split | (Kos *et al.*, 2015) |
| AZPAE14968 | 2009 | Croatia | Split | (Kos *et al.*, 2015) |
| AZPAE14969 | 2010 | United States | Roseburg | (Kos *et al.*, 2015) |
| AZPAE14970 | 2010 | United States | Roseburg | (Kos *et al.*, 2015) |
| AZPAE14971 | 2010 | China | Shatin | (Kos *et al.*, 2015) |
| AZPAE14972 | 2010 | Germany | Aachen | (Kos *et al.*, 2015) |
| AZPAE14973 | 2010 | Germany | Aachen | (Kos *et al.*, 2015) |
| AZPAE14974 | 2010 | Germany | Aachen | (Kos *et al.*, 2015) |
| AZPAE14975 | 2010 | China | Beijing | (Kos *et al.*, 2015) |
| AZPAE14976 | 2010 | China | Beijing | (Kos *et al.*, 2015) |
| AZPAE14977 | 2010 | China | Beijing | (Kos *et al.*, 2015) |
| AZPAE14978 | 2010 | United States | Roseburg | (Kos *et al.*, 2015) |
| AZPAE14979 | 2010 | United States | Roseburg | (Kos *et al.*, 2015) |
| AZPAE14980 | 2010 | United States | Detroit | (Kos *et al.*, 2015) |
| AZPAE14981 | 2010 | France | Paris | (Kos *et al.*, 2015) |

Continued on next page

**Table S1 – continued from previous page**

| Isolat | Year of isolation | Country | City | Reference |
|---|---|---|---|---|
| AZPAE14982 | 2010 | Croatia | Split | (Kos *et al.*, 2015) |
| AZPAE14983 | 2010 | Croatia | Split | (Kos *et al.*, 2015) |
| AZPAE14984 | 2010 | France | Paris | (Kos *et al.*, 2015) |
| AZPAE14985 | 2010 | China | Shatin | (Kos *et al.*, 2015) |
| AZPAE14986 | 2010 | China | Shatin | (Kos *et al.*, 2015) |
| AZPAE14987 | 2010 | Germany | Koln | (Kos *et al.*, 2015) |
| AZPAE14988 | 2010 | China | Shatin | (Kos *et al.*, 2015) |
| AZPAE14989 | 2010 | China | Hong Kong | (Kos *et al.*, 2015) |
| AZPAE14990 | 2010 | China | Hong Kong | (Kos *et al.*, 2015) |
| AZPAE14991 | 2010 | Spain | Santander | (Kos *et al.*, 2015) |
| AZPAE14992 | 2010 | Germany | München | (Kos *et al.*, 2015) |
| AZPAE14993 | 2010 | Spain | Madrid | (Kos *et al.*, 2015) |
| AZPAE14994 | 2010 | Germany | Heidelberg | (Kos *et al.*, 2015) |
| AZPAE14995 | 2010 | Spain | Palma de Mallorca | (Kos *et al.*, 2015) |
| AZPAE14996 | 2010 | Spain | Palma de Mallorca | (Kos *et al.*, 2015) |
| AZPAE14997 | 2010 | Spain | Palma de Mallorca | (Kos *et al.*, 2015) |
| AZPAE14998 | 2010 | Spain | Palma de Mallorca | (Kos *et al.*, 2015) |
| AZPAE14999 | 2010 | Spain | Bilbao | (Kos *et al.*, 2015) |
| AZPAE15000 | 2010 | Spain | Bilbao | (Kos *et al.*, 2015) |
| AZPAE15001 | 2011 | Colombia | Bogota | (Kos *et al.*, 2015) |
| AZPAE15002 | 2010 | Spain | Bilbao | (Kos *et al.*, 2015) |
| AZPAE15003 | 2011 | Colombia | Bogota | (Kos *et al.*, 2015) |
| AZPAE15004 | 2011 | Colombia | Bogota | (Kos *et al.*, 2015) |
| AZPAE15005 | 2011 | United States | Roseburg | (Kos *et al.*, 2015) |
| AZPAE15006 | 2011 | United States | Roseburg | (Kos *et al.*, 2015) |
| AZPAE15007 | 2010 | Spain | Santander | (Kos *et al.*, 2015) |
| AZPAE15008 | 2010 | Spain | Santander | (Kos *et al.*, 2015) |
| AZPAE15009 | 2010 | Spain | Santander | (Kos *et al.*, 2015) |
| AZPAE15010 | 2010 | Spain | Santander | (Kos *et al.*, 2015) |
| AZPAE15011 | 2010 | Spain | Santander | (Kos *et al.*, 2015) |
| AZPAE15012 | 2011 | Germany | Koln | (Kos *et al.*, 2015) |
| AZPAE15013 | 2011 | Germany | Koln | (Kos *et al.*, 2015) |
| AZPAE15014 | 2011 | Germany | Koln | (Kos *et al.*, 2015) |
| AZPAE15015 | 2011 | Germany | Koln | (Kos *et al.*, 2015) |
| AZPAE15016 | 2011 | United States | Detroit | (Kos *et al.*, 2015) |
| AZPAE15017 | 2011 | United States | Detroit | (Kos *et al.*, 2015) |
| AZPAE15018 | 2011 | United States | Detroit | (Kos *et al.*, 2015) |
| AZPAE15019 | 2010 | France | Besancon | (Kos *et al.*, 2015) |

Continued on next page

**Table S1 – continued from previous page**

| Isolat | Year of isolation | Country | City | Reference |
|---|---|---|---|---|
| AZPAE15020 | 2010 | France | Besancon | (Kos *et al.*, 2015) |
| AZPAE15021 | 2011 | Argentina | Victoria | (Kos *et al.*, 2015) |
| AZPAE15022 | 2011 | France | Nantes | (Kos *et al.*, 2015) |
| AZPAE15023 | 2011 | Spain | Madrid | (Kos *et al.*, 2015) |
| AZPAE15024 | 2011 | Spain | Madrid | (Kos *et al.*, 2015) |
| AZPAE15025 | 2011 | Spain | Madrid | (Kos *et al.*, 2015) |
| AZPAE15026 | 2011 | Colombia | Bogota | (Kos *et al.*, 2015) |
| AZPAE15027 | 2011 | Spain | Bilbao | (Kos *et al.*, 2015) |
| AZPAE15028 | 2011 | France | Paris | (Kos *et al.*, 2015) |
| AZPAE15029 | 2011 | France | Paris | (Kos *et al.*, 2015) |
| AZPAE15030 | 2011 | Germany | München | (Kos *et al.*, 2015) |
| AZPAE15031 | 2011 | Germany | Aachen | (Kos *et al.*, 2015) |
| AZPAE15032 | 2011 | France | Rouen | (Kos *et al.*, 2015) |
| AZPAE15033 | 2011 | France | Rouen | (Kos *et al.*, 2015) |
| AZPAE15034 | 2011 | Spain | Bilbao | (Kos *et al.*, 2015) |
| AZPAE15035 | 2011 | Spain | Bilbao | (Kos *et al.*, 2015) |
| AZPAE15036 | 2012 | China | Shatin | (Kos *et al.*, 2015) |
| AZPAE15037 | 2012 | France | Paris | (Kos *et al.*, 2015) |
| AZPAE15038 | 2012 | France | Paris | (Kos *et al.*, 2015) |
| AZPAE15039 | 2012 | Germany | Heidelberg | (Kos *et al.*, 2015) |
| AZPAE15040 | 2012 | Germany | Heidelberg | (Kos *et al.*, 2015) |
| AZPAE15041 | 2012 | Germany | Koln | (Kos *et al.*, 2015) |
| AZPAE15042 | 2012 | Germany | Heidelberg | (Kos *et al.*, 2015) |
| AZPAE15043 | 2012 | France | Nantes | (Kos *et al.*, 2015) |
| AZPAE15044 | 2012 | France | Nantes | (Kos *et al.*, 2015) |
| AZPAE15045 | 2011 | France | Nantes | (Kos *et al.*, 2015) |
| AZPAE15046 | 2012 | Argentina | Victoria | (Kos *et al.*, 2015) |
| AZPAE15047 | 2012 | Argentina | Victoria | (Kos *et al.*, 2015) |
| AZPAE15048 | 2012 | Germany | München | (Kos *et al.*, 2015) |
| AZPAE15049 | 2012 | Germany | München | (Kos *et al.*, 2015) |
| AZPAE15050 | 2012 | China | Hong Kong | (Kos *et al.*, 2015) |
| AZPAE15051 | 2012 | China | Hong Kong | (Kos *et al.*, 2015) |
| AZPAE15052 | 2012 | Argentina | Victoria | (Kos *et al.*, 2015) |
| AZPAE15053 | 2012 | Argentina | Victoria | (Kos *et al.*, 2015) |
| AZPAE15054 | 2012 | Colombia | Bogota | (Kos *et al.*, 2015) |
| AZPAE15055 | 2012 | Colombia | Bogota | (Kos *et al.*, 2015) |
| AZPAE15056 | 2012 | China | Beijing | (Kos *et al.*, 2015) |
| AZPAE15057 | 2012 | China | Beijing | (Kos *et al.*, 2015) |

Continued on next page

**Table S1 – continued from previous page**

| Isolat | Year of isolation | Country | City | Reference |
|---|---|---|---|---|
| AZPAE15058 | 2012 | France | Nantes | (Kos *et al.*, 2015) |
| AZPAE15059 | 2012 | France | Nantes | (Kos *et al.*, 2015) |
| AZPAE15060 | 2012 | France | Nantes | (Kos *et al.*, 2015) |
| AZPAE15061 | 2012 | France | Nantes | (Kos *et al.*, 2015) |
| AZPAE15062 | 2012 | Brazil | Curitiba | (Kos *et al.*, 2015) |
| AZPAE15063 | 2012 | Brazil | Curitiba | (Kos *et al.*, 2015) |
| AZPAE15064 | 2012 | Brazil | Curitiba | (Kos *et al.*, 2015) |
| AZPAE15065 | 2012 | Brazil | Curitiba | (Kos *et al.*, 2015) |
| AZPAE15066 | 2003 | Croatia | Split | (Kos *et al.*, 2015) |
| AZPAE15067 | 2004 | Germany | Heidelberg | (Kos *et al.*, 2015) |
| AZPAE15068 | 2004 | Germany | Heidelberg | (Kos *et al.*, 2015) |
| AZPAE15069 | 2004 | Germany | Heidelberg | (Kos *et al.*, 2015) |
| AZPAE15070 | 2004 | Germany | Heidelberg | (Kos *et al.*, 2015) |
| AZPAE15071 | 2004 | Germany | Heidelberg | (Kos *et al.*, 2015) |
| AZPAE15072 | 2004 | Germany | Heidelberg | (Kos *et al.*, 2015) |
| PAO1 | 1954 | Australia | Melbourne | (Holloway, 1955) |
| PA14 | Unknown | Unknown | Unknown | (Rahme *et al.*, 1995) |
| PA7 | Unknown | Argentina | Unknown | (Roy *et al.*, 2010) |

**Table S2. Primer sequences used to construct mutant genotypes.** The forward inner primers (those named with inner For) include a single capital letter (in bold) which coded for the site directed mutation of interest. Lower case letters represent sequence homology or spacers, the un-bolded capital letters represent restriction enzyme (Res. Enz.) recognition sequence.

| Name | Primer Sequence | Res. Enz. |
|---|---|---|
| gyrA_inner_For | gcgGGTCTCgcgaca**T**cgcggtctacgacaccatc | BsaI |
| gyrA_outer_Rev | gcgGGTCTCAGCAAgccaccacgttgatgccg | BsaI |
| gyrA_inner_Rev | gcgGGTCTCtgtcgccgtgcgggtgg | BsaI |
| gyrA_outer_For | gcgGGTCTCAGTCGgcgaggacatcccgatcgaag | BsaI |
| parC_outer_For | gcgGAAGACaCAATTGACTAGTatcatcccctaaccagcgcc | BbsI, MfeI, SpeI |
| parC_inner_Rev | gcgGAAGACcggcctgctacgaggcc | BbsI |
| parC_S87L_inner_For | gcgGAAGACgcaggcc**A**agtcgccgtgcgggtgg | BbsI |
| parC_S87W_inner_For | gcgGAAGACgcaggcc**C**agtcgccgtgcgggtgg | BbsI |
| parC_outer_Rev | gcgGAAGACGGCGCGCCtactacgccctcgacgaagc | BbsI, AscI |

**Table S3. Correlation between know resistance determining loci and resistance phenotype.** With knowledge of the resistance phenotype for strains in our alignment, we counted which strains had mutations in the codon of amino acid (AA) positions known to confer fluoroquinolone resistance. We used a chi-squared test for the independence between mutation at these positions and the resistance phenotype and highlight significant ($P \leq 0.05$) values in bold. Recall there were 389 strains in our alignment of which 192 (197) were susceptible (resistant).

| Gene | AA | Mutants | Resistant | $X^2$ | df | P |
|---|---|---|---|---|---|---|
| *gyrA* | 83 | 161 | 156 | 231.941 | 1 | **2.250** $\times 10^{-52}$ |
| | 87 | 34 | 33 | 30.110 | 1 | **4.083** $\times 10^{-08}$ |
| *gyrB* | 466 | 16 | 14 | 7.596 | 1 | **0.006** |
| | 468 | 9 | 9 | 7.072 | 1 | **0.008** |
| *parC* | 87 | 128 | 127 | 177.208 | 1 | **1.979** $\times 10^{-40}$ |
| | 91 | 6 | 6 | 4.103 | 1 | **0.043** |
| *parE* | 457 | 8 | 7 | 3.061 | 1 | 0.080 |
| | 459 | 4 | 3 | 0.227 | 1 | 0.634 |
| | 473 | 11 | 7 | 0.323 | 1 | 0.570 |
| *morA* | 563 | 14 | 7 | 0.000 | 1 | 1.000 |
| | 975 | 15 | 6 | 0.333 | 1 | 0.564 |
| | 1056 | 15 | 6 | 0.333 | 1 | 0.564 |
| | 1109 | 15 | 7 | 0.003 | 1 | 0.960 |
| | 1155 | 16 | 9 | 0.041 | 1 | 0.839 |
| | 1162 | 15 | 8 | 0.000 | 1 | 1.000 |
| | 1213 | 15 | 7 | 0.003 | 1 | 0.960 |

**Table S4. Evidence of relative $\Delta G$ epistasis among the strongly correlated pairs of synonymous intragenic substitutions.** Epistasis is measured with a multiplicative model and error is calculated using error propagation (Trindade *et al.*, 2009). There is evidence for epistasis when the absolute value of is greater than the error of our measures. We find no evidence for epistasis as $\epsilon$ is never greater than estimates of error.

| **Mutant Pair** | | $\epsilon$ | **Error** |
|---|---|---|---|
| *dnaN* c495t | *dnaN* c504t | $-9.263 \times 10^{-4}$ | $2.387 \times 10^{-2}$ |
| *gyrB* c1422t | *gyrB* c1443t | $-1.481 \times 10^{-4}$ | $4.768 \times 10^{-3}$ |
| *morA* a4041g | *morA* c4083t | $1.113 \times 10^{-2}$ | $3.394 \times 10^{-2}$ |
| *parC* c1533t | *parC* c1581t | $-7.982 \times 10^{-3}$ | $2.794 \times 10^{-2}$ |
| *parC* c1533t | *parC* c1587t | $8.039 \times 10^{-4}$ | $2.514 \times 10^{-2}$ |
| *parC* c1533t | *parC* t1554g | $-1.798 \times 10^{-2}$ | $3.166 \times 10^{-2}$ |
| *parC* c1581t | *parC* c1587t | $4.633 \times 10^{-4}$ | $2.635 \times 10^{-2}$ |
| *parC* t1554g | *parC* c1581t | $-6.985 \times 10^{-3}$ | $3.529 \times 10^{-2}$ |

**Table S5. Evidence of $I_{TE}$ epistasis among the strongly correlated pairs of synonymous intragenic substitutions.** Epistasis is measured with a multiplicative model and error is calculated using error propagation (Trindade *et al.*, 2009). There is evidence for epistasis when the absolute value of is greater than the error of our measures. We find negative epistasis for the sites in *morA* based on $\epsilon$ having an absolute value greater than the estimates of error.

| Mutational Pair | | $\epsilon$ | Error |
|---|---|---|---|
| *dnaN* c504t | *dnaN* c495t | $1.4113 \times 10^{-5}$ | $3.234 \times 10^{-3}$ |
| *gyrB* c1443t | *gyrB* c1422t | $5.330 \times 10^{-5}$ | $1.334 \times 10^{-2}$ |
| *morA* c4083t | *morA* a4041g | $\mathbf{-4.437} \times 10^{-2}$ | $\mathbf{1.053} \times 10^{-2}$ |
| *parC* c1581t | *parC* c1587t | $1.390 \times 10^{-5}$ | $4.606 \times 10^{-3}$ |
| *parC* c1533t | *parC* c1587t | $1.362 \times 10^{-5}$ | $7.304 \times 10^{-3}$ |
| *parC* t1554g | *parC* c1581t | $1.769 \times 10^{-6}$ | $2.770 \times 10^{-3}$ |
| *parC* c1533t | *parC* c1581t | $3.148 \times 10^{-5}$ | $6.041 \times 10^{-3}$ |
| *parC* c1533t | *parC* t1554g | $-1.025 \times 10^{-5}$ | $5.360 \times 10^{-3}$ |

**Table S6. Test for independence between the relative position of correlated substitutions and the type of pair they form.** We used a $\chi^2$ test and present the observed (expected) counts for the number of significantly correlated pairs ($P \leq 10^{-4}$) of substitutions based on if they were in the same gene and how many of the substitutions were synonymous. Analysis led to a test statistic of 7246.04 with 2 degrees of freedom and $P \sim 0$.

| Within the same gene | Type of Pair | | |
|---|---|---|---|
| | **Non Synonymous** | **One Synonymous** | **Synonymous** |
| **TRUE** | 0<br>(0.52) | 6<br>(1.83) | 60<br>(1.63) |
| **FALSE** | 101<br>(1289.31) | 2274<br>(4569.33) | 7470<br>(4048.38) |

**Table S7. Absorbance (600 nm) readings from ciprofloxacin MIC assay of *P. aeruginosa* WT and mutant constructs.** Readings were taken after 24 hours of growth on a 96 well plate where wells contained half salt LB growth medium with ciprofloxacin concentration as denoted by the column heading. The values presented are the mean (standard deviation) values for a minimum of 4 replicates. No genotype appeared more than once on a single 96 well plate.

| | | Concentration of Ciprofloxacin ($Log_2\ \mu g/mL$) | | | | | | | | | | | |
|---|---|---|---|---|---|---|---|---|---|---|---|---|---|
| Background | Mutation | 5 | 4 | 3 | 2 | 1 | 0 | -1 | -2 | -3 | -4 | -5 | -6 |
| PA01 | WT | 0.01 | 0.021 | 0.02 | 0.012 | 0.127 | 0.021 | 0.04 | 0.326 | 0.417 | 0.536 | 0.675 | 0.916 |
| | | (0.006) | (0.004) | (0.002) | (0.001) | (0.156) | (0.004) | (0.02) | (0.034) | (0.001) | (0.074) | (0.273) | (0.006) |
| | *gyrA* c248t | 0.013 | 0.017 | 0.1 | 0.398 | 0.521 | 0.512 | 0.754 | 0.764 | 0.772 | 0.779 | 0.814 | 0.912 |
| | | (0.004) | (0.007) | (0.042) | (0.149) | (0.087) | (0.029) | (0.134) | (0.087) | (0.078) | (0.091) | (0.068) | (0.09) |
| | *parC* c260t | 0.016 | 0.031 | 0.036 | 0.014 | 0.053 | 0.034 | 0.059 | 0.447 | 0.468 | 0.504 | 0.639 | 0.894 |
| | | (0.003) | (0.017) | (0.034) | (0.006) | (0.057) | (0.025) | (0.019) | (0.03) | (0.037) | (0.052) | (0.06) | (0.122) |
| | *parC* c260g | 0.015 | 0.034 | 0.022 | 0.018 | 0.014 | 0.029 | 0.098 | 0.422 | 0.463 | 0.499 | 0.698 | 0.859 |
| | | (0.005) | (0.018) | (0.006) | (0.001) | (0.002) | (0.019) | (0.114) | (0.057) | (0.053) | (0.06) | (0.063) | (0.129) |
| | *gyrA* c248t, | 0.418 | 0.76 | 0.843 | 0.803 | 0.845 | 0.805 | 0.754 | 0.748 | 0.686 | 0.807 | 0.774 | 0.854 |
| | *parC* c260t | (0.116) | (0.094) | (0.138) | (0.113) | (0.172) | (0.13) | (0.145) | (0.138) | (0.138) | (0.085) | (0.091) | (0.139) |
| | *gyrA* c248t, | 0.376 | 0.598 | 0.629 | 0.768 | 0.814 | 0.836 | 0.918 | 0.81 | 0.826 | 0.854 | 0.881 | 0.982 |
| | *parC* c260g | (0.386) | (0.179) | (0.116) | (0.132) | (0.057) | (0.07) | (0.059) | (0.099) | (0.111) | (0.038) | (0.033) | (0.071) |
| PA14 | WT | 0.044 | 0.054 | 0.063 | 0.084 | 0.094 | 0.1 | 0.02 | 0.02 | 0.04 | 0.285 | 0.615 | 0.936 |
| | | (0.006) | (0.005) | (0.01) | (0.012) | (0.017) | (0.093) | (0.004) | (0.003) | (0.019) | (0.22) | (0.163) | (0.086) |
| | *gyrA* c248t | 0.07 | 0.079 | 0.087 | 0.314 | 0.25 | 0.133 | 0.264 | 0.864 | 0.895 | 0.952 | 0.899 | 1.096 |
| | | (0.013) | (0.005) | (0.036) | (0.071) | (0.034) | (0.05) | (0.145) | (0.122) | (0.054) | (0.059) | (0.065) | (0.035) |
| | *parC* c260t | 0.04 | 0.064 | 0.096 | 0.12 | 0.114 | 0.07 | 0.022 | 0.033 | 0.025 | 0.226 | 0.659 | 0.819 |
| | | (0.007) | (0.004) | (0.008) | (0.028) | (0.018) | (0.013) | (0.003) | (0.013) | (0.003) | (0.085) | (0.119) | (0.169) |
| | *parC* c260g | 0.045 | 0.095 | 0.1 | 0.13 | 0.121 | 0.07 | 0.028 | 0.04 | 0.03 | 0.249 | 0.649 | 0.961 |
| | | (0.004) | (0.025) | (0.058) | (0.06) | (0.047) | (0.04) | (0.006) | (0.003) | (0.004) | (0.086) | (0.1) | (0.145) |
| | *gyrA* c248t, | 0.279 | 0.294 | 0.237 | 0.769 | 0.996 | 0.957 | 0.941 | 0.956 | 0.933 | 0.948 | 0.927 | 0.941 |
| | *parC* c260t | (0.012) | (0.029) | (0.038) | (0.184) | (0.077) | (0.056) | (0.019) | (0.018) | (0.027) | (0.028) | (0.018) | (0.012) |
| | *gyrA* c248t, | 0.195 | 0.25 | 0.268 | 0.809 | 1.054 | 0.956 | 0.92 | 0.926 | 0.921 | 0.921 | 0.932 | 0.987 |
| | *parC* c260g | (0.078) | (0.041) | (0.096) | (0.064) | (0.073) | (0.048) | (0.056) | (0.022) | (0.035) | (0.025) | (0.038) | (0.125) |

### SI Figures

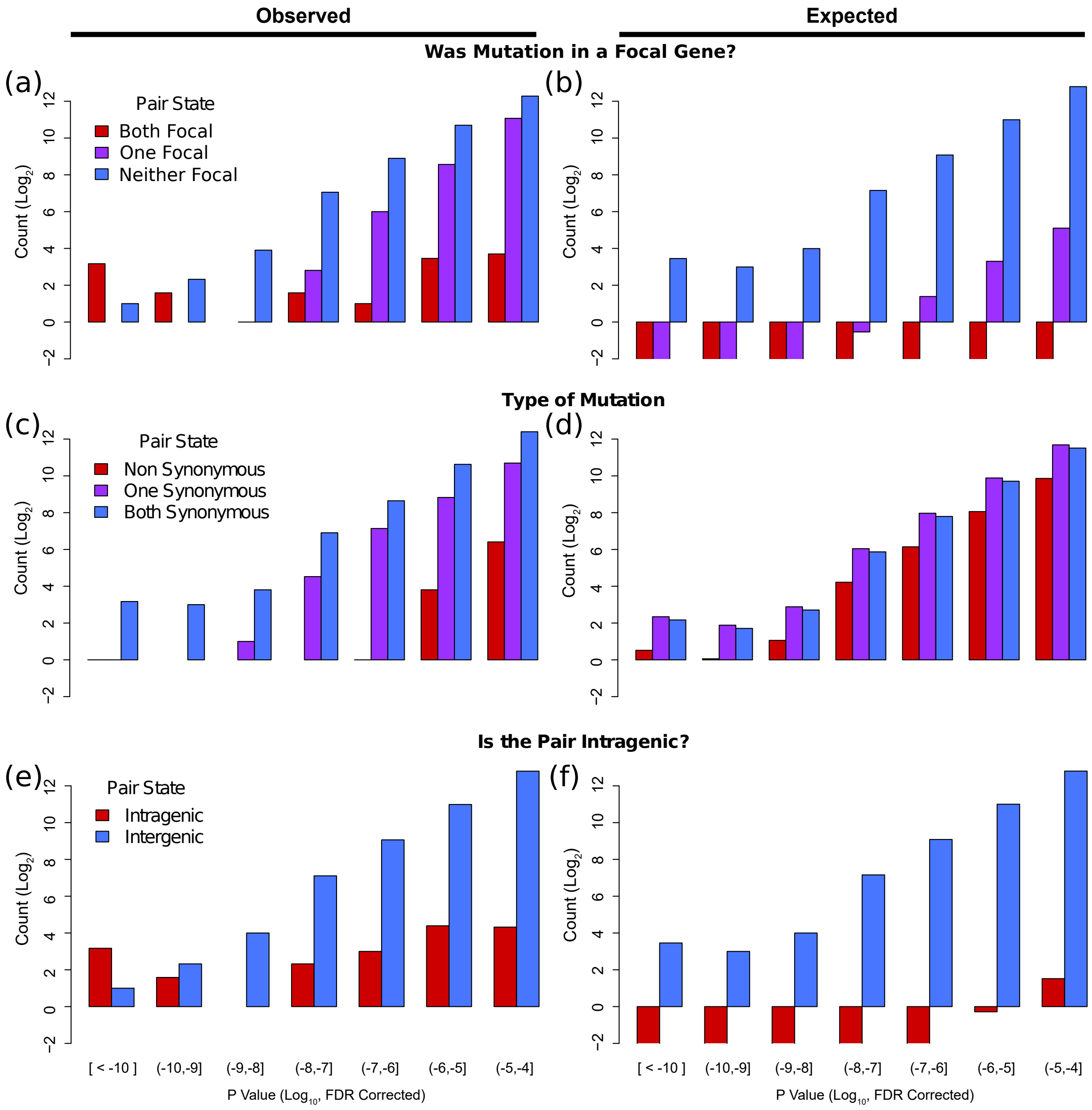


**Figure S1. Comparisons of observed and expected types of correlated substitutions with at least weak ($P \leq 10^{-4}$) support.** Note that all y-axes are based on $\log_2$ counts and so when there is no bar the count was 0. Panels a,c,e show the observed distributions of when correlated pairs were: (a) in genes we expected to show correlation, (c) nonsynonymous, or (e) intragenic. Whereas b,d,f show the null expected distributions, of their left-most counterpart, if a same number of correlated pairs had been randomly drawn from our alignment.

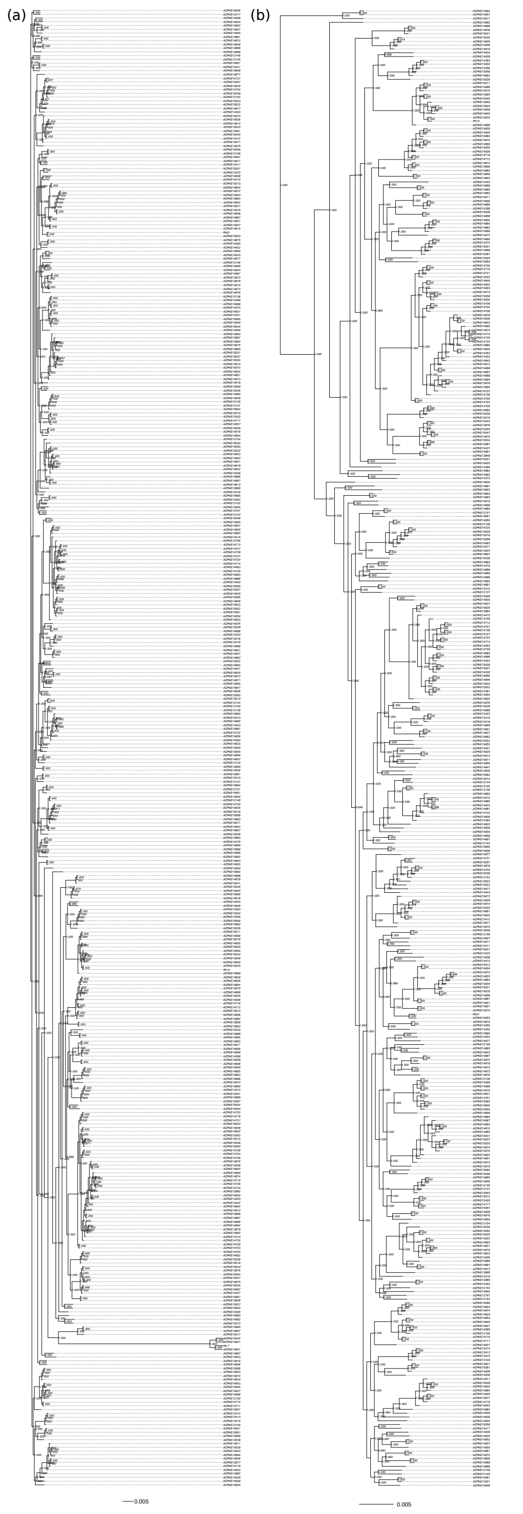


**Figure S2. Phylogenetic trees estimated from the concatenated alignment of all conserved Information class, determined from COG terms, genes in our dataset.** Support for bifurcations are printed at nodes and were estimated using FastTree's default SH test for local support values. (a) shows the unrooted tree which includes the taxonomic outliers identified by being part of the long branch sub-clade including PA7. (b) shows the rooted tree once the PA7 sub-clade was removed. Analysis of correlated evolution, with AEGIS, was performed with the tree presented in (b).

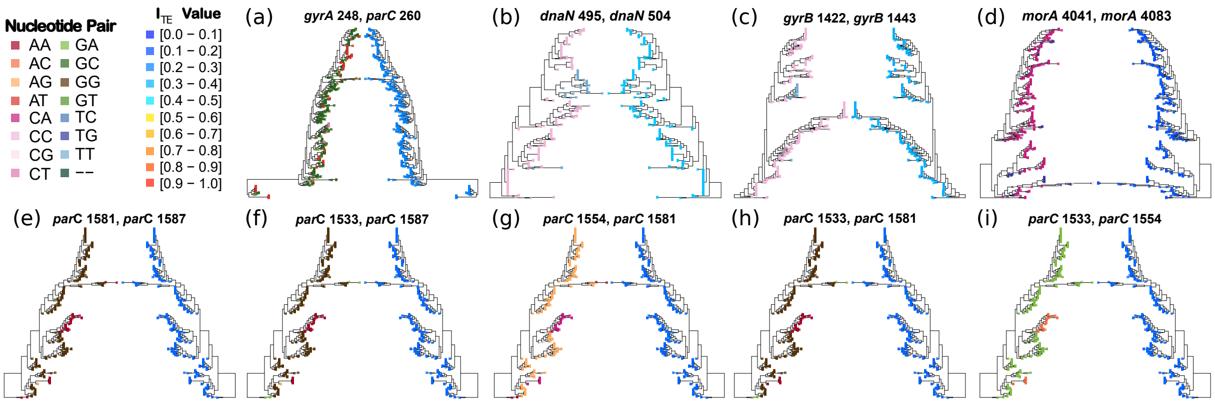


**Figure S3. Phylogeny of the gene(s) which contain the most significantly correlated pairs of substitutions ($P < 10^{11}$).** Each tree was constructed with FastTree (Price *et al.*, 2010) (GTR $+\Gamma_5$). The title of each panel indicates the pair of substitutions being considered. For trees on the left hand side, the color of the tips represent the paired nucleotide state of strains. On the right side, the color of tips represent the $I_{TE}$ value of the whole gene(s) sequence.

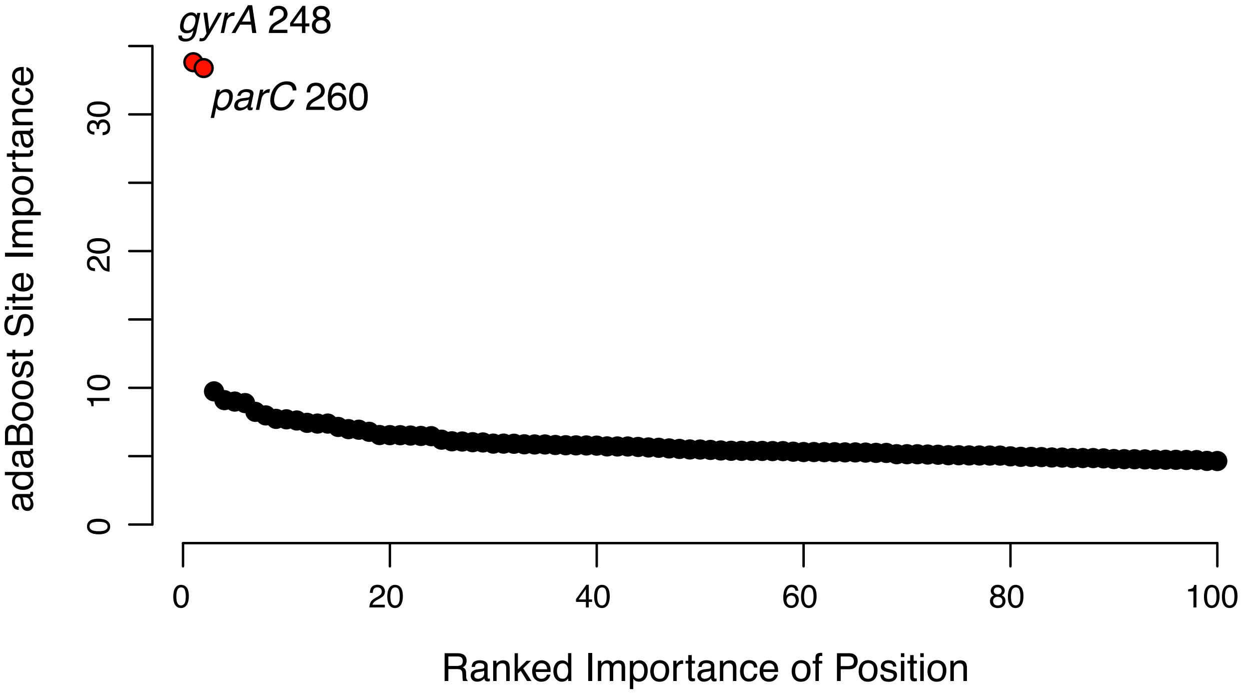


**Figure S4. Top 100 ranked importance values in predicting levofloxacin (a fluoroquinolone drug) resistance for nucleotide positions in our alignment.** The importance values were obtained by running the adaptive boosting machine learning algorithm *boosting* (implemented in the R package *adabag* (Alfaro *et al.*, 2013)) on all polymorphic sites in our alignment. Levofloxacin resistance phenotype information was obtained from previously published data (Kos *et al.*, 2015). Importance values reflect the strength of correlation between genomic sites and the resistance phenotype. Red circles denote genomic positions which have been previously reported in the literature to correlate with fluoroquinolone resistance.

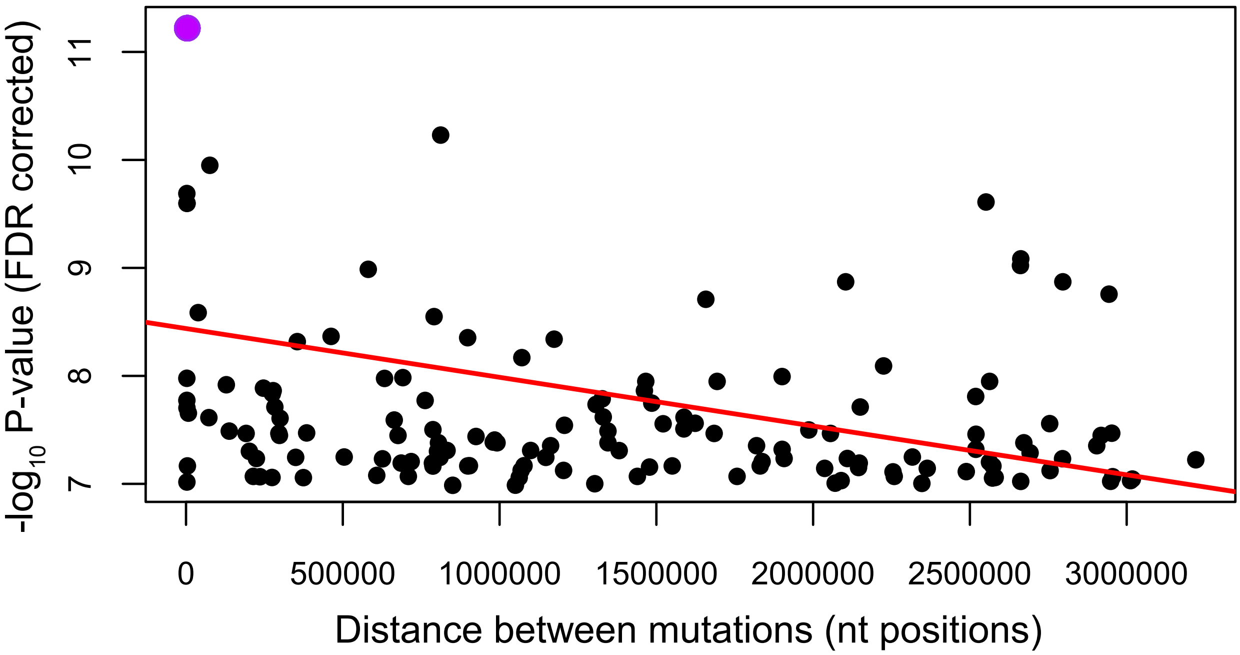


**Figure S5. The signal of correlated evolution for paired synonymous substitutions as a function of physical distance.** Physical distance is measured as the number of base pairs, using the circular reference chromosome of PA14, separating two mutations. The red line shows the linear regression for all significantly correlated pairs with at least medium support ($P \leq 10^{-7}$). Purple highlights the most significantly correlated pairs of substitutions.
